# Habitat complexity alters the strength of sexual selection on brain size in a livebearing fish

**DOI:** 10.1101/2025.03.20.644301

**Authors:** Léa Daupagne, Alessandro Devigili, Rebecca McNeil, David Wheatcroft, Niclas Kolm, John L. Fitzpatrick

**Affiliations:** Department of Zoology, Stockholm University, Svante Arrhenius väg 18B, Stockholm, 10691, Sweden; Department of Biology, University of Padova, Padova, 35131, Italy

**Keywords:** Mate choice, Sexual selection, Habitat complexity, Halfbeaks, Brain size

## Abstract

Animals often reproduce in complex environments, which should generate selection for both enhanced detectability in signaling traits and improved cognitive processing abilities. However, the extent to which signaling and cognitive traits have evolved to overcome the challenges of interacting in complex habitats remains understudied. We examined whether habitat complexity influences sexual selection in the pygmy halfbeak, *Dermogenys collettei*, a small livebearing freshwater fish. Using free-swimming arenas, we created low- and high-complexity environments and observed mating behaviors in mixed-sex groups. While the opportunity for sexual selection did not differ significantly between environments for either sex, we observed positive selection gradients for both male and female brain size in open arenas, but not in complex habitats. Interestingly, no clear selection on male or female ornamentation was detected in either environment. Our findings suggest that habitat complexity may limit selection pressures on cognitive traits, such as brain size, without significantly affecting visual ornamentation’s role in mating success.

## 1 Introduction

The strength and direction of sexual selection can vary among populations (McLain, 1982; Prohl, 2002; Kasumovic et al., 2008), and both within (Wacker et al., 2014; Carleial et al., 2023; Daupagne et al., 2023) and between breeding cycles (Madsen and Shine, 1993; Cockburn et al., 2008; Siepielski et al., 2009). Such spatio-temporal variation in sexual selection is expected to be a powerful mechanism for maintaining genetic diversity in sexually selected traits (Chaine and Lyon, 2008), making it a central focus in evolutionary biology. When the relative performance of different alleles is dependent on the environment in which they are expressed, genotype-by-environment interactions can prevent the loss of genetic variation in sexual traits (Hunt and Hosken, 2014; Ellner and Hairston Jr, 1994). This spatio-temporal variation in the strength of sexual selection may also affect the sexes differently (e.g. Singh and Punzalan, 2018). Males typically experience stronger sexual selection than females, likely driven by underlying differences in gametic investment (i.e., anisogamy, Janicke et al., 2016; Davies et al., 2023; Janicke, 2024). Consequently, variation in the strength of sexual selection across time and space may be more extreme in males than females. However, identifying the specific environmental and social factors that drive this variation in selection remains challenging. While much of the research on sexual selection draws on long-term observational studies in natural populations, such studies often struggle to disentangle the co-varying ecological factors influencing selection (Cornwallis and Uller, 2010). For instance, studies of wild populations indicate that sexual selection on male traits can be influenced by complex interactions between abiotic factors (e.g., temperature and humidity, Le Galliard et al., 2005) and between biotic and demographic factors (e.g., predation pressure and population density Endler, 1980; McKellar et al., 2009; Arendt et al., 2014; Reznick et al., 2001). To better understand these dynamics, controlled experimental studies are needed to tease apart the effects of these interacting factors on sexual selection.

An ecological factor of particular interest is habitat complexity (i.e., the structural heterogeneity of an environment), as it has important effects on behavioral processes related to sexual selection (e.g., Moyaho et al. 2004; van der Sluijs et al. 2011; Myhre et al. 2013). Detecting and discerning sexual signals become more challenging in complex habitats, as the efficacy of sexual signals displayed by courters and evaluated by choosers is influenced by the physical environment (Endler, 1992). For example, in habitats with dense canopy cover, animals experience limited transmission of visual signals over long distances due to reduced light availability. This can either decrease their mating interactions (e.g., Endler 1993; Thery 2001; Hibler and Houde 2006; Devigili et al. 2021) or increase courtship complexity and effort (e.g., Gomes et al. 2017; Miles and Fuxjager 2018), thereby altering selection pressures on courter sexual traits (e.g., Endler and Basolo 1998; Boughman 2002). Similarly, habitat complexity can affect intra-sexual competition and selection on traits linked to aggression and competitive success (Hibler and Houde, 2006; Lackey and Boughman, 2013). For instance, male-male competition in sticklebacks (*Gasterosteus aculeatus*) is higher in open than in complex habitats, which might generate stronger selection for larger body size (Lackey and Boughman, 2013). Similarly, in male-dimorphic mites (*Rhizoglyphus echinopus*), fewer fighter males were produced in populations evolving in complex environments compared to open environments, demonstrating that habitat complexity can alter the strength of selection on pre-copulatory male-male competitive traits (Tomkins et al., 2011). Although habitat complexity can influence sexual and antagonistic behaviors by altering conspecific encounter rates and affecting demography, less attention has been given to how habitat complexity directly impacts variance in reproductive success and the strength of selection on male and female sexual traits. Understanding these dynamics is crucial for comprehending how environmental factors shape evolutionary processes.

Habitat complexity can also exert selection on brain morphology. At the macro-evolutionary scale, specific brain regions are larger in species that live in more complex habitats (e.g., echolocating bats, Safi and Dechmann, 2005; chipmunks, Budeau and Verts, 1986; and Tanganyikan cichlids, Shumway, 2008). Similar patterns are present at the within-species level; for example, hippocampus volumes are larger in Leach’s storm petrels (*Oceanodroma leucorhoa*) nesting in complex wooded habitats compared to open meadow habitats (Abbott et al., 1999). Complex environments are thought to be more cognitively demanding, requiring animals to navigate a greater number of potential paths to locate food and shelter, thus selecting for increased neural investment. Larger brains are generally associated with increased cognitive abilities (Striedter, 2005), and recent experimental evidence demonstrates that variation in brain size is directly related to cognitive performance on a range of tasks (e.g., Kotrschal et al., 2013, 2014*a*, 2015; Buechel et al., 2018). Importantly, habitat complexity may also exert selection on brain size due to the links between the physical environment and processes related to sexual selection. The perceptual and cognitive processes associated with sexual selection (i.e., mate identification, evaluation, comparison and choice) are intimately linked with the brain (Ryan, 2021). For instance, Corral-Ĺopez et al. (2017) demonstrated that large-brained female guppies (*Poecilia reticulata*) are better at assessing attractive males than small-brained females, which highlights the link between brain size, cognitive ability and mate choice (Culumber et al., 2020; McNeil et al., 2021). Yet, the interaction between brain size, sexual selection and habitat complexity remains unclear. If complex habitats make it more difficult to detect and discern differences in sexual signals, then investing more in neural resources might improve fitness in complex habitats by allowing individuals to make more accurate mating decisions. Alternatively, if increased neural investment does not fully mitigate the challenges of detecting and comparing mates in complex habitats, the energetic costs of a larger brain may outweigh its cognitive benefits for mate choice, leading to a negative correlation between brain size and habitat complexity. Furthermore, the strength and direction of selection on neural tissue could differ between courters and choosers, depending on how brain size influences fitness in these respective mating roles. Clarifying how habitat complexity influences selection on brain size thus requires experimental studies that examine both courters and choosers.

In this study, we investigate how habitat complexity influences the strength and direction of sexual selection in the pygmy halfbeak (*Dermogenys collettei*), a small viviparous fish native to Southeast Asia. Pygmy halfbeaks are an excellent model for studying the impact of habitat complexity on sexual behaviors, as they are found in a wide range of habitats including streams, rivers, lakes and estuaries of varying habitat complexities, both around (e.g., canopy cover) and within (e.g., filamentous algae, aquatic vegetation) their aquatic environments (Ng and Tan, 1999; Baker and Lim, 2012; Devigili et al., 2021). In addition, halfbeaks live in mixed-sex groups where intra- and inter-sexual interactions are common (Greven et al., 2010; Devigili et al., 2021). Halfbeaks frequently engage in agonistic interactions (Reuland et al., 2021; Devigili et al., 2021), which can escalate to ‘wrestling’ matches where individuals interlock their elongated lower jaws (called beaks) and attempt to displace one another in often prolonged contest (Berten and Greven, 1991; Greven, 2006; Greven et al., 2010). Halfbeaks also exhibit sexual dimorphism in body size, with females being larger than males, and sexual dichromatism, with males displaying more red and yellow coloration, particularly on their modified anal fin, the andropodium (Greven et al., 2010; Reuland et al., 2020). Female mate choice is influenced by the amount of red coloration displayed by males, albeit inconsistently (Reuland et al., 2020; McNeil et al., 2021), while male mate choice is influenced by the size of a sex-specific orange gravid spot on the female’s abdomen that varies in size with the reproductive cycle (Ogden et al., 2020). Importantly, unlike other well studied internally fertilizing fishes (e.g., Poeciliids, Magurran and Seghers, 1994; Evans, 2010), pygmy halfbeaks do not engage in sneak matings, which suggests that males are not able to undermine female preferences and that pre-copulatory interactions are an important route to fitness in these fish. Pygmy halfbeaks have semi-transparent heads, allowing for noninvasive external estimates of the size of individual brain areas (McNeil et al., 2021), which enables experimental exploration of the relationship between brain size variation and sexual behaviors in environments with varying complexity.

Here, we quantify and compare the strength and direction of selection on brain size, body size, beak size and sexual coloration in both male and female halfbeaks interacting in experimental arenas where habitat complexity was either low (open arenas, Figure 1A) or complex (spatially partitioned arenas, Figure 1B). Since there is reduced scope for visual and behavioral interactions between individuals in the complex areas compared to the open arenas, we generated a series of predictions on how habitat complexity might influence the strength and direction of sexual selection. First, we predict that, regardless of habitat complexity, the strength of sexual selection will be stronger in males than females (Janicke et al., 2016; Fromhage and Jennions, 2016; Janicke and Morrow, 2018). Second, we predict that selection on brain size will be stronger in complex environments, where - assuming a link between brain size and cognition - processing and responding to social information, such as tracking interactions with potential mates and competitors over time, may be critical. Third, we predict that competitive interactions will be less common in complex arenas (Tomkins et al., 2011), and therefore selection on body and beak size will be stronger in open compared to complex environments. We also expect that the stronger selection on body and beak size will be more pronounced in males than females, as agonistic interactions are more common among male halfbeaks (Devigili et al., 2021). Finally, we predict that complex arenas will dampen mate choice by preventing simultaneous comparison of potential mates (i.e., sequential mate choice can reduce the scope for mate choice, Barry and Kokko, 2010). Since both male and female halfbeaks engage in mate choice (Ogden et al., 2020; Reuland et al., 2021), we expect that male sexual ornaments (i.e., red and yellow coloration), female gravid spot size, and body size (in both sexes) will be under stronger selection in open than complex arenas.

**Fig. 1:**
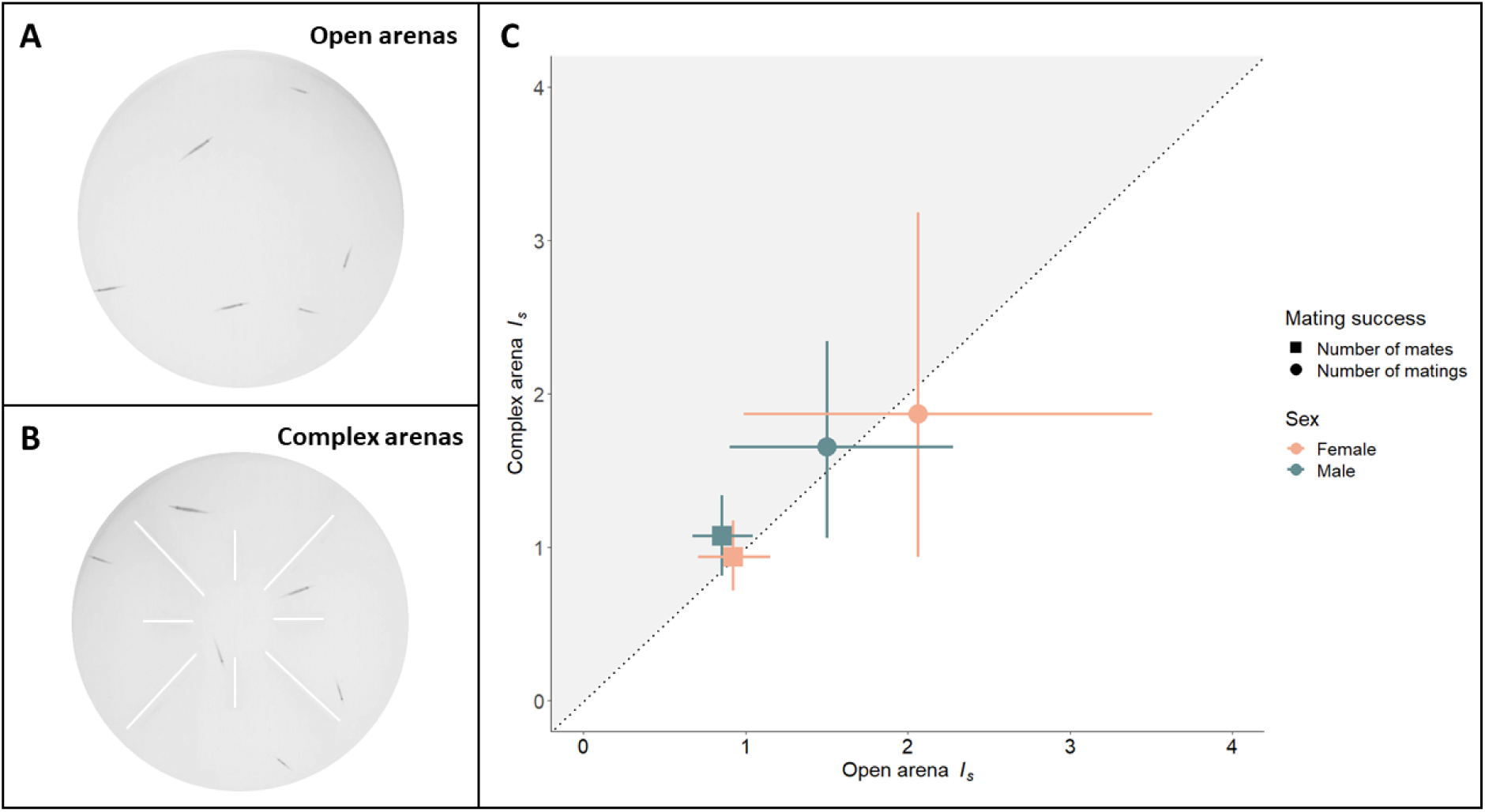
Habitat complexity and the opportunity for sexual selection. **A**: Representation of an *open* arena. Eight opaque barriers, four measuring 10 cm and three measuring 5 cm, were present within the arena. **B**: Representation of a *complex* arena. Fish could interact in an unobstructed environment. **C**: Opportunity for sexual selection (*I_s_*) estimated in males (blue) and females (orange) in complex (*y* axis) or open arenas (*x* axis) for both mating success characterisation (*MS_M_*: number of matings; *MS_P_*: number of mates). Mean estimates and 95% CI are obtained from bootstrapping (Table 2). Dotted line mark equal effect size for both arena types. Grey area above the diagonal indicates the space in which *I_s_* is found to be larger in the complex arena. The *x/y* distance of mean estimates from dashed line is equal to the effect size (Table 2).

## 2 Methods

### 2.1 Study population and rearing conditions

Experimental fish were descendants of wild-caught halfbeaks sourced from the Tebrau River, Malaysia, that were housed in mix-sex stock tanks in an aquarium facility of # University using a standard set of husbandry practices. Briefly, all aquaria were oxygenated and contained 2 cm of gravel and artificial plants. A 12:12 dark/light cycle was maintained within the laboratory and the temperature was kept at 27 ± 1.5 °C. Fish were fed a mix of flake food and recently hatched *Artemia salina* nauplii (artemia; Anostraca: Artemiidae). To produce fish for the experiment, gravid females were removed from stock tanks and placed in 7.5 L tanks where they were monitored daily. Upon giving birth, offspring were separated from mothers to prevent maternal cannibalism and kept in 5 L tanks with up to five other fry. The onset of sexual maturity was determined by identifying maturing males by their thickened andropodium, a modified anal fin used for sperm transfer (Greven et al., 2010). At the onset of sexual maturity, males and females were placed into sex-specific tanks.

### 2.2 Experimental design

Reproductive behaviors were quantified in sexually mature fish housed in circular mating arenas made of white plastic (35 cm in diameter, 4 cm water depth, Figure 1A,B). Fish were randomly allocated to one of two experimental arena types. In the *open* arenas (n=37), fish could interact in an unobstructed environment (Figure 1A). In the *complex* arenas (n=37), we increased the complexity of the spatial environment by adding eight opaque barriers, four measuring 10 cm and three measuring 5 cm, within the arena (Figure 1B). Each trial involved three males and three females, maintaining a 1:1 sex ratio. Fish were initially placed in individually labeled opaque containers inside the mating arenas for a 15 min habituation period, during which the fish had the opportunity to recover from being placed in a new environment. These labels, along with morphological differences, enabled individual identification during behavior scoring (see below). After the habituation period, the opaque containers were lifted, allowing fish to interact for 7 hours. Mating behaviors were recorded at 60 frames per second with a camera (Point Grey Grasshopper 3 4.1 megapixel camera with Fujinon CF25HA-1 lens) placed 1-5 m above the mating arenas. Five trials were excluded from the dataset due to escaped individuals or absence of matings, resulting in 69 trials in total (34 in open arenas and 35 in complex arenas).

### 2.3 Calculating mating success

Using individual labels for each fish, mating behaviors of every individual were recorded throughout the 7 hour period where fish could interact freely. Specifically, we focused on quantifying the number of matings performed during the trials. Halfbeaks are internal fertilizers, with males courting females by swimming under them and then attempting to mate by rapidly (∼ 40-80 ms) twisting their body in an attempt to use their andropodium to transfer sperm to females (Greven et al., 2010). Because halfbeaks are specialized surface dwellers, the rapid body movement creates ripples in the water. This makes mating a conspicuous, albeit brief, behavior, that can be observed from the cameras mounted above the mating arenas. For each presumed mating, we re-examined the behavior frame-by-frame and only recorded a mating attempt as having occurred when the male’s body was twisted into a characteristic C-shape. Whenever copulations were observed, we recorded the identity of each individual involved. This allowed us to calculate two sex-specific metrics to quantify the variance in mating success among individuals of the same sex: i) the number of matings performed by the focal individual with all its mates (*MS_M_*) and ii) the number of partners with which the focal individual mated (*MS_P_*). *MS_M_* and *MS_p_* were calculated separately for males and females.

### 2.4 Quantifying phenotypic traits

Following interactions in the mating arena, body size and external traits were quantified using digital photographs. Specifically, we measured body length, beak size, brain size, male coloration (red and yellow), and female gravid spot size, traits that may be targeted by sexual selection. Body length is a common target of mate choice (Andersson, 1994; Andersson and Iwasa, 1996, reviewed by Hunt et al., 2009) and an indicator of fish condition (i.e., their fecundity, Wooton, 1979; Morita and Takashima, 1998 and quality, Reynolds and Gross, 1992). In halfbeaks, beaks are weapons used in intrasexual competition (Greven et al., 2010). Brain size, although not directly observable by conspecifics due to the surface-dwelling ecology of halfbeaks, may influence mating success through its role in cognitive processes involved in mate choice and competition (Corral-Ĺopez et al., 2017). Red and yellow coloration was quantified in males as these traits have been hypothesized to serve as a sexual ornament in halfbeaks (Coleman et al. 2009; red only: Reuland et al. 2020). Finally, gravid spot size, an orange abdominal marking that varies in size among females and across the female reproductive cycle, was quantified in females as gravid spot size is used by males to make mate choice decisions (Ogden et al., 2020).

For males and females, the lateral side of each fish was photographed using a Canon 800D digital camera fitted with a macro LF-S 60 mm lens inside a photo chamber (30 x 20 x 20 cm) filled with water under standard light conditions. Each image included a scale bar to facilitate subsequent measurements. For each individual, we measured body length (from the anterior part of the eye to the caudal peduncle) (mm) and beak length (from the anterior tip of the beak to the anterior part of the eye) (mm) using Image J v1.52i (Schneider et al., 2012). Although body length and beak length both exhibit condition dependent expression in halfbeaks (Fernlund Isaksson et al., 2022), we considered these phenotypic traits separately as males are more likely to engage in intra-sexual competition than females (Devigili et al., 2021) and body size may experience different selective pressures than beak size between the sexes (e.g., body size may be selected for in intra-sexual selection in males and be an indicator of fecundity used in inter-sexual sexual selection in females). Using the polygon selection tool in ImageJ, we also measured the area of red and yellow coloration on males body and fins (mm^2^) and the gravid spot area (mm^2^) in females.

We then placed males and females in a clear plastic photo chamber (7.5 x 5 x 2.5 cm) filled with water and used a Canon 800D digitial camera to take a photograph of the dorsal surface of each individual. Each photograph included a scale bar. Halfbeaks have semi-transparent heads that allow internal structures of the brain to be measured in a non-invasive manner (McNeil et al., 2021). From these dorsal photos we measured the total dorsal brain area (mm^2^), which consists of the dorsal portion of the olfactory bulbs, telencephalon, optic tectum, cerebellum and dorsal medula, using ImageJ (McNeil et al., 2021). Our previous work demonstrated that there is a significant pairwise repeatability between total external dorsal brain area and measures of total brain volume and weight from dissected brains (McNeil et al., 2021). Therefore, our external measure of total dorsal brain area is a good proxy for total neural investment.

### 2.5 Statistical analysis

Statistical analyses were carried out using R version 4.3.0 (Team, 2023). We used complementary metrics of sexual selection (Table 1) to estimate and to compare the strength of pre-copulatory sexual selection in each sex and for each arena type.

**Table 1:** Different metrics of sexual selection used in this study.

| Measure | Description | Interpretation |
| --- | --- | --- |
| Opportunity for sexual selection ($I_s$) | Standardized variance in mating success | Upper strength of sexual selection |
| Mating differentials ($m'$) | Covariance between relative mating success and the standardized trait values | Direction selection operating on a single trait |
| Linear selection gradients ($\beta$) | Vector of directional selection coefficients ($\beta$) obtained from multiple linear regressions of relative mating success on standardized traits | Directional selection operating on multiple traits |
| Non-linear selection gradients ($\gamma$) | Matrix of quadratic and correlational selection coefficients ($\gamma$) obtained from multiple linear regressions of relative mating success on standardized traits | Stabilizing or disruptive selection operating on multiple traits |

**Table 2:** Influence of arena type on the opportunity of sexual selection (*I_s_*).

| Parameter | Sex | Mean complex<br>(95% CI) | Mean open<br>(95% CI) | Effect size<br>(95% CI) | *P-values* |
| --- | --- | --- | --- | --- | --- |
| $MS_M$ | Female | 1.99 (0.82,3.57) | 2.21 (0.96,3.89) | -0.22 (-2.28,1.85) | 0.82 |
|  | Male | 1.72 (0.98,2.55) | 1.50 (0.80,2.39) | 0.23 (-0.93,1.34) | 0.68 |
| | $\Delta\text{Sex}$ | -0.25 (-1.68,0.96)<br>*p-value*=0.72 | -0.56 (-2.11,0.78)<br>*p-value*=0.44 | - | - |
| $MS_P$ | Female | 0.92 (0.68, 1.18) | 0.96 (0.70, 1.23) | -0.03 (-2.32, 1.82) | 0.87 |
|  | Male | 1.10 (0.78, 1.43) | 0.83 (0.65, 1.04) | 0.28 (-0.10, 0.65) | 0.16 |
| | $\Delta\text{Sex}$ | 0.14 (-0.21,0.48)<br>*p-value*=0.44 | -0.07 (-0.36,0.21)<br>*p-value*=0.63 | - | - |

Confidence intervals in each arena type are obtained via bootstrap sampling and are shown in each sex and for both mating success characterization ( $MS_M$ : number of matings; $MS_P$ : number of mates). Sex difference observed within each arena type ( $\Delta\text{Sex}$ ) are also represented. Significance of effects sizes is assessed via permutations tests (*P-values*). Significance effects are shown in boldface.

#### 2.5.1 Opportunity for sexual selection (*I_s_*)

The opportunity for sexual selection (*I_s_*) is a standardized selection metric measured in each sex as the variance in mating success (here *MS_M_*: number of matings and *MS_P_*: number of mates) divided by the square of the mean mating success (Wade, 1979; Wade and Arnold, 1980). We first calculated *I_s_* in each arena and performed (1) a two-way ANOVA to compare *I_s_* between sexes and arena types (i.e., open vs. complex) and (2) a one-way ANOVA within each sex to compare *I_s_* between arena types. An interaction term was initially included in the two-way ANOVA models, but was removed as non-significant. We then calculated *I_s_* at the scale of the arena type, i.e., considering all arenas of one type as a simple group. We used bootstrapping with 10,000 bootstrap replicates (R package ‘boot’; Canty and Ripley 2017; Davison and Hinkley 1997) to estimate variances with their 95% confidence intervals (95% CI). *I_s_* were compared between arena types and between sexes by computing pairwise differences of bootstrap samples and their 95% CIs. More specifically, arena type comparisons were done by subtracting the open arenas estimates from the complex arena estimates while sex comparisons were done by subtracting the female estimates from the male estimates. Such differences reflect the effect size; negative differences indicate larger values in the open arenas or in females while positive differences indicate larger values in the complex arenas or in males. Two-sided permutation tests (10,000 permutations) were additionally performed to access whether this difference (i.e., the effect size) is statistically significant.

#### 2.5.2 Mating differentials (*m’*), linear (*β*) and non-linear (*γ*) selection gradients

In addition to adopting a variance based approach, we used trait-based measures of sexual selection, including mating differentials (*m’*), linear selection gradients (*β*) and non-linear selection gradients (*γ*) on traits that are likely under selection. Selection on standard body length, beak length and brain area was investigated in both sexes. In males, we additionally investigated selection on the amount of red and yellow coloration present on their body and fins. To obtain standardized metrics of sexual selection, we first relativized mating success (both *MS_M_* and *MS_P_*) within each sex and arena type by dividing each observed value by the mean value of that sex and arena type. We then standardized each trait by subtracting the mean trait value and by dividing by the standard deviation of the trait (Jones, 2009).

Mating differentials (*m’*) were calculated at the arena level as the covariance between the standardized trait values and relative mating success (Lande and Arnold, 1983). In the same way that is described above for *I_s_*, we used a bootstrapping approach to obtain covariances with their 95% confidence intervals and two sided-permutations tests to compare covariances values between arena types and between sexes. We also performed mating differentials on relative morphological traits by selecting the residuals from the regression of each trait (e.g., brain size) on body length (Table S13).

Linear selection gradients (*β*) and the matrix of non-linear (quadratic and correlational) selection gradients (*γ*), were estimated using a multiple regression approach for each sex and each arena type. We estimated *β*, which indicates directional selection, with linear regressions and *γ* with a model incorporating all linear, quadratic, and cross-product terms. Quadratic regression coefficients were doubled to obtain estimates of non-linear (stabilizing/disruptive) selection gradients (Stinchcombe et al., 2008). The estimation of the strength of non-linear selection based on the quadratic coefficients alone can be biased as it does not include the effect of correlational selection on paired traits (Phillips and Arnold, 1989; Blows and Brooks, 2003). We therefore investigated non-linear selection by performing a canonical rotation of the *γ* matrix to find the major axes of the fitness surface (Phillips and Arnold, 1989; Blows and Brooks, 2003). Each new axis is described by an eigenvector *m* in which trait representation is similar to that of a principal component analysis. The form of non-linear selection (stabilizing/disruptive) along each eigenvector *m* is indicated by the sign of the associated eigenvalue (*λ*) (positive = disruptive selection; negative = stabilizing selection), while the strength of selection (how curve the surface is) is given by its size. To assess the significance of non-linear selection on each of the eigenvectors obtained from canonical rotation of the fitness surface, we performed a standard permutation procedure (10,000 permutations) as recommended by Reynolds et al. (2010), using the R script provided by the authors in the supplementary material.

In females, selection gradients were performed on (1) beak length, (2) standard length, (3) brain area and (4) gravid spot area, while in males gradients were performed on (1) beak length, (2) standard length, (3) brain area, (4) total red area and (5) total yellow area. We measured the amount of multicollinearity in the regression analysis by using a variance inflation factor (VIF). As multicollinearity was detected in females’ models, we additionally performed a model excluding the variable with the highest VIF value (i.e., standard length) to improve the accuracy and stability of the regression analysis. These models exhibited similar statistical outputs and are presented in supplementary (Tables S9, S10, S11, and S12).

### Ethical statement

All experiments were conducted in accordance with the Animal Research Ethical Board (permit number 2393-2018). These applications are consistent with the Institutional Animal Care and Use Committee guidelines.

## 3 Results

### 3.1 General description of the mating behaviors

A total of 202 (±1.40, 0-10, ±SD, range) and 213 (±1.38, 0-10) matings were observed in open and complex arenas, respectively. On average, 5.9 (±4.4, 1-22) matings occurred in open arenas, while 6.1 (±4.7, 1-27) matings occured in complex arenas. For males, the average number of mating partners was 0.91 (±0.85, 0-3) in open arenas and 0.92 (±0.95, 0-3) in complex arenas. Similarly, for females, the average number of mating partners was 0.91 (±0.88, 0-3) in open arenas and 0.92 (±0.89, 0-3) in complex arenas.

### 3.2 Opportunity for sexual selection (*I_s_*)

First, we calculated the opportunity for sexual selection (*I_s_*) in each arena and compared *I_s_* between arena types and sexes. There was no effect of arena type or sex on *I_s_* for *MS_M_* (ANOVA: F_1_,_134_ = 0.18, P = 0.67, and F_1_,_134_ = 0.005, P = 0.94, respectively). Similarly, we observed no effect of arena type or sex on *I_s_* for *MS_P_* (ANOVA: F_1_,_134_ = 0.25, P = 0.62, and F_1_,_134_ = 0.006, P = 0.84, respectively). We also compared *I_s_* between arena types within each sex. In females, there was no effect of arena type on *I_s_* either for *MS_M_* or *MS_P_* (ANOVA: F_1_,_67_ = 0, P = 0.991, and F_1_,_67_ = 0.007, P = 0.935, respectively). Similarly, there was no effect of arena type on *I_s_* in males, either for *MS_M_* or *MS_P_* (ANOVA: F_1_,_67_ = 0.37, P = 0.545, and F_1_,_67_ = 0.39, P = 0.535, respectively, Figure S1). We then calculated *I_s_* at the scale of the arena type and used bootstrapping to estimate variances with their 95% confidence intervals (95% CI). There was no effect of arena type on *I_s_* in both sexes, either for *MS_P_* or *MS_M_* (Table 2 and Figure 1). Similarly, there was no significant sex difference in *I_s_* within each arena type (ΔSex), either for *MS_P_* or *MS_M_*.

### 3.3 Mating differentials (*m’*), linear (*β*) and non-linear (*γ*) selection gradients

#### 3.3.1 Mating differentials (*m’*)

When mating success was calculated as the number of matings (*MS**_M_***), mating differentials (*m’*) on beak length and standard length in females were significantly positive in open arenas, but not in complex arenas (Table 3 and Figure S1). These findings indicate that there is positive directional selection on female beak length and standard length in open arenas. *m’* on female brain area was significantly positive in both open and complex arenas (Table 3 and Figure S1), indicating positive directional selection on female brain area in both arena types. In males, there was no mating differentials significantly different from zero in either arena type (Table 3 and Figure S2). We detected sex differences in *m’* on beak length, standard length and brain area in open arenas, but not in complex arenas, with females having larger *m’* compared to males (Table 3).

**Table 3:** Influence of arena type on mating differentials (*m’*) on morphological traits.

| Parameter | Mean complex<br>(95% CI) | Mean open (95%<br>CI) | Effect size (95%<br>CI) | *P-values* |
| --- | --- | --- | --- | --- |
| *MS<sub>M</sub>* |  |  |  |  |
| **Females** |  |  |  |  |
| Beak length | 0.12 (-0.08,0.32) | **0.43 (0.21,0.68)** | -0.31 (-0.63,-0.002) | 0.11 |
| Standard length | 0.16 (-0.04,0.35) | **0.45 (0.25,0.66)** | -0.29 (-0.58,-0.01) | 0.13 |
| Brain area | **0.25 (0.05,0.47)** | **0.50 (0.30,0.72)** | -0.25(-0.54,0.04) | 0.19 |
| Gravid spot area | 0.07 (-0.16,0.31) | 0.20 (-0.07,0.45) | -0.12 (-0.48,0.24) | 0.51 |
| **Males** |  |  |  |  |
| Beak length | -0.10 (-0.34,0.12) | -0.13 (-0.34,0.07) | 0.03(-0.28,0.32) | 0.86 |
| Standard length | -0.11 (-0.34,0.11) | -0.14 (-0.38,0.09) | 0.03 (-0.29,0.35) | 0.85 |
| Brain area | -0.05 (-0.28,0.18) | -0.06 (-0.27,0.15) | 0.00 (-0.30, 0.32) | 0.98 |
| Total red areas | 0.10 (-0.15,0.36) | 0.02 (-0.22,0.28) | 0.08 (-0.28,0.43) | 0.63 |
| Total yellow areas | -0.13 (-0.39,0.09) | -0.12 (-0.33,0.08) | -0.01 (-0.34,0.29) | 0.93 |
| **ΔSex** |  |  |  |  |
| Beak length | -0.22 (-0.53,0.07)<br>*p-value* = 0.22 | **-0.56 (-0.89,-0.26)**<br>*p-value* = **0.003** | - | - |
| Standard length | -0.27 (-0.56,0.04)<br>*p-value* = 0.14 | **-0.59 (-0.91,-0.27)**<br>*p-value* = **0.002** | - | - |
| Brain area | -0.31 (-0.62,0.00)<br>*p-value* = 0.09 | **-0.60 (-0.87,-0.27)**<br>*p-value* = **0.003** | - | - |
| *MS<sub>P</sub>* |  |  |  |  |
| **Females** |  |  |  |  |
| Beak length | 0.11 (-0.06,0.29) | **0.33 (0.15,0.51)** | -0.22 (-0.47,0.03) | 0.10 |
| Standard length | 0.15 (-0.03,0.33) | **0.38 (0.21,0.56)** | -0.23 (-0.48,0.01) | 0.07 |
| Brain area | **0.21 (0.04,0.37)** | **0.40 (0.23,0.59)** | -0.19(-0.45,0.05) | 0.14 |
| Gravid spot area | 0.12 (-0.07,0.30) | **0.27 (0.08,0.49)** | -0.16 (-0.44,0.12) | 0.24 |
| **Males** |  |  |  |  |
| Beak length | -0.14 (-0.35,0.06) | -0.12 (-0.30,0.06) | -0.02(-0.30,0.25) | 0.88 |
| Standard length | -0.06 (-0.24,0.11) | -0.11 (-0.30,0.09) | 0.04 (-0.22,0.31) | 0.75 |
| Brain area | 0.03 (-0.15,0.21) | 0.00 (-0.18,0.17) | 0.03 (-0.22, 0.28) | 0.82 |
| Total red areas | 0.05 (-0.13,0.23) | 0.04 (-0.16,0.24) | 0.01 (-0.26,0.28) | 0.94 |
| Total yellow areas | -0.02 (-0.21,0.17) | -0.15 (-0.32,0.02) | 0.13 (-0.12,0.38) | 0.32 |
| **ΔSex** |  |  |  |  |
| Beak length | -0.25 (-0.52,0.01)<br>*p-value* = 0.07 | **-0.45 (-0.70,0.19)**<br>*p-value* = **0.001** | - | - |
| Standard length | -0.21 (-0.46,0.04)<br>*p-value* = 0.13 | **-0.49 (-0.75,0.23)**<br>*p-value* < **0.001** | - | - |
| Brain area | -0.18 (-0.43,0.07)<br>*p-value* = 0.19 | **-0.41 (-0.66,-0.15)**<br>*p-value* = **0.002** | - | - |

Confidence intervals in each arena type are obtained via bootstrap sampling and are shown for both sexes, for all traits and for both mating success characterisation ( $MS_M$ : number of matings; $MS_P$ : number of mates). Sex difference observed within each arena type ( ΔSex) are also represented. Significance of effects sizes is assessed via permutations tests (*P-values*). Significance effects are shown in boldface.

**Table 4:**
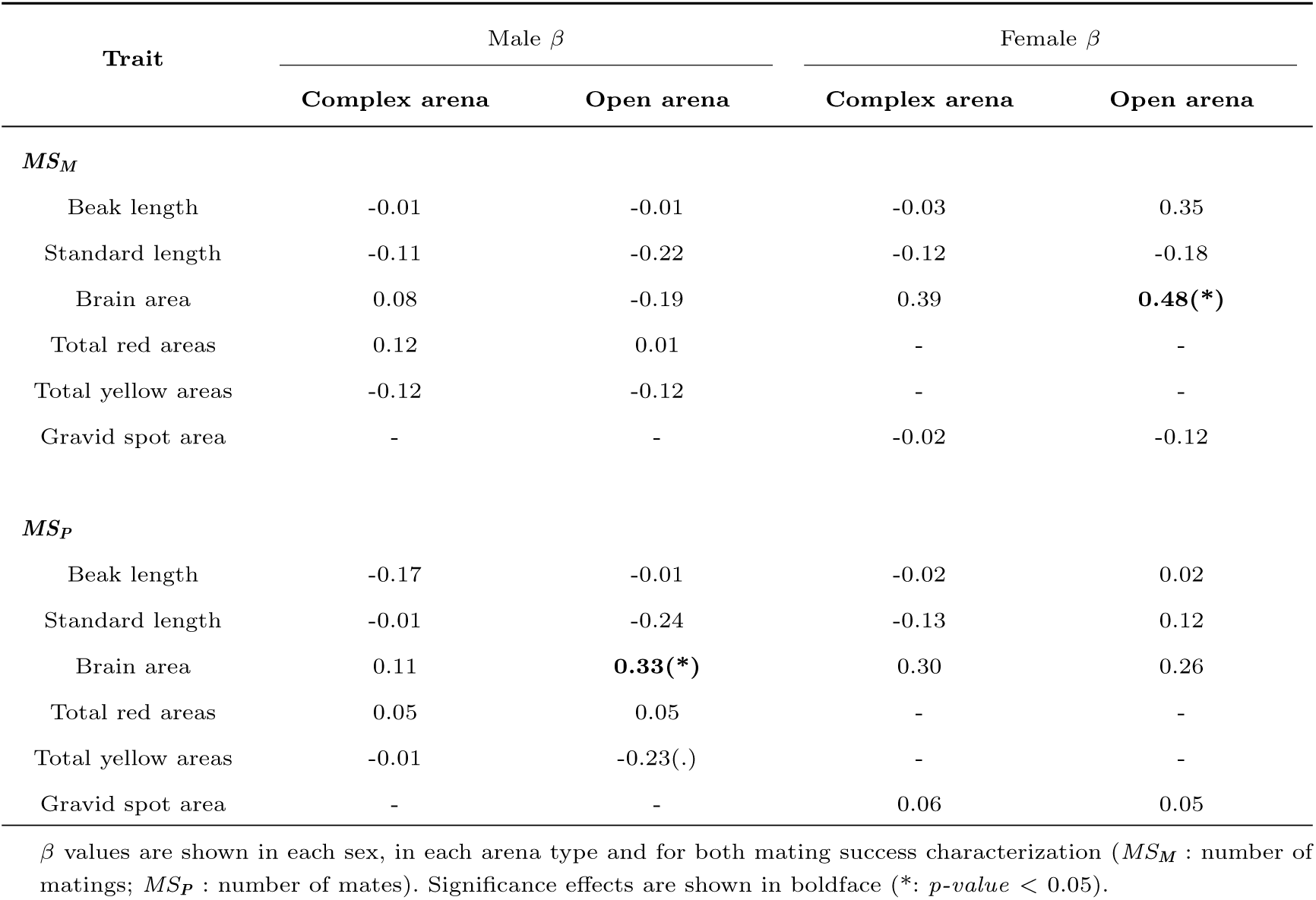
Linear (*β*) selection gradients on morphological traits.

| Trait | Male $\beta$ | | Female $\beta$ | |
| --- | --- | --- | --- | --- |
|  | Complex arena | Open arena | Complex arena | Open arena |
| **$MS_M$** |  |  |  |  |
| Beak length | -0.01 | -0.01 | -0.03 | 0.35 |
| Standard length | -0.11 | -0.22 | -0.12 | -0.18 |
| Brain area | 0.08 | -0.19 | 0.39 | **0.48(\*)** |
| Total red areas | 0.12 | 0.01 | - | - |
| Total yellow areas | -0.12 | -0.12 | - | - |
| Gravid spot area | - | - | -0.02 | -0.12 |
| **$MS_P$** |  |  |  |  |
| Beak length | -0.17 | -0.01 | -0.02 | 0.02 |
| Standard length | -0.01 | -0.24 | -0.13 | 0.12 |
| Brain area | 0.11 | **0.33(\*)** | 0.30 | 0.26 |
| Total red areas | 0.05 | 0.05 | - | - |
| Total yellow areas | -0.01 | -0.23(.) | - | - |
| Gravid spot area | - | - | 0.06 | 0.05 |

$\beta$ values are shown in each sex, in each arena type and for both mating success characterization ( $MS_M$ : number of matings; $MS_P$ : number of mates). Significance effects are shown in boldface (\*: $p$ -value < 0.05).

When mating success was calculated as the number of partners (*MS**_P_***), *m’* on beak length, standard length and brain area in females were similar to those obtained for *MS**_M_*** (Table 3). Additionally, *m’* on gravid spot area was significantly positive in open arenas, but not in complex arenas (Table 3). No mating differentials significantly differed from zeros in males in either arena type (Table 3). Females in open arenas had larger *m’* in beak length, standard length and brain area compared to males (Table 3). In contrast, *m’* did not differ between the sexes in complex arenas (Table 3).

Overall, there was no significant influence of arena type on *m’* for any of the traits investigated, suggesting that arena complexity does not impose differential selection pressure on each individual trait examined. Moreover, when controlling for body size, only relative brain area showed consistent positive selection across arena types and mating success characterisation in females (Table S12), and no significant sex differences were detected in relative trait values.

#### 3.3.2 Linear (*β*) and non-linear (*γ*) selection gradients

We detected significant positive linear (*β*) selection on brain area size in open arenas in both females (with relative mating success calculated as the number of matings (*MS**_M_***), Tables 4 and S1) and males (with relative mating success calculated as the number of partners (*MS**_P_***), Tables 4 and S2). This indicates that large brained females have a higher number of mating partners while large brained males mate at a higher rate. There was no evidence of linear (*β*) selection in open arenas for any other trait examined in either females or males (Tables 4, S1 and S2). In complex arenas, there was no evidence of linear (*β*) selection on any of the examined traits in females or males (Tables 4, S3 and S4).

In the open arenas, there was no evidence of non-linear selection (*γ*) on any of the traits examined in either females (Tables S1 and S5) or males (Tables S2 and S6). However, following canonical rotation of the *γ* matrix we detected stabilizing selection on eigenvector *m3* in males in complex arenas (Table S4), with relative mating success calculated as *MS**_M_*** (Table S8). The *m3* vector was primarily loaded by total red area. This pattern suggests that males with intermediate area of red coloration mate at a higher rate. There was no evidence of non-linear selection on *m* vectors for males in open arenas (Tables S2 and S6) or for females in either open (Tables S1 and S5) or complex arena (Tables S3 and S7).

## 4 Discussion

Our study offers novel insights into how habitat complexity shapes sexual selection dynamics. By experimentally quantifying the strength and direction of selection on a range of traits in pygmy halfbeaks, we identified key targets of selection, as well as traits that are broadly unaffected by the imposed degree of habitat complexity. Brain size emerged as a key factor influencing mating success, primarily in open environments. This suggests that open environments lead to strong selection on cognitive processes linked with fitness, a finding that challenges the conventional expectation that larger brains provide fitness advantages in more complex habitats. The opportunity for sexual selection (*I_s_*) did not differ between arena types for either sex, aligning with the general pattern observed across the animal kingdom (Janicke et al., 2016) and consistent with species exhibiting polygynandrous mating systems (Hare and Simmons, 2019). However, despite similar *I_s_* values, we detected sex-specific selection, with evidence of stronger selection on female beak, body and brain size than male traits, but only in open environments. Surprisingly, we found little evidence that sexual ornaments potentially involved in mate choice (i.e., male coloration in males and female gravid spots) were under selection in our experimental set-up. These findings suggest that increasing habitat complexity can limit selection pressures on cognitive traits linked with fitness without simultaneously affecting visual ornamentation which is commonly assumed to be an integral component of mating success.

Greater investment in brain tissue increased fitness in both female and male halfbeaks. However, the relationship between brain size and fitness differed between open and complex environments. In open environments, females with larger brains successfully mated more often, while males with larger brains mated with a greater number of partners. These sex-specific patterns associated with brain size suggest different routes to increase fitness between females and males. For example, increased mating frequency in females may reflect greater proactivity in securing mating opportunities, which could enhance fertilization success. In contrast, in males, mating with a greater number of partners generally provides a more direct route to increased fitness through higher reproductive output (Bateman, 1948). Regardless of the route to increase fitness, our findings suggest a strong link between brain size and fitness in halfbeaks. The lack of evidence that selection acts on brain size in complex environments was surprising. We initially hypothesized that larger brains would provide greater advantages in more complex environments, where restricted movement and limited visibility increase the need for enhanced cognitive abilities, such as problem-solving and adaptability (e.g., Brown and Warburton, 1997). For example, studies on cichlids have demonstrated that visual acuity and spatial memory are enhanced in populations inhabiting more complex environments (Dobberfuhl et al., 2005). Furthermore, comparative studies across multiple species, as well as within-species analysis, have consistently reported a positive correlation between brain size and habitat complexity in nature (Ratcliffe et al., 2006; Safi and Dechmann, 2005). However, in complex habitats, selection pressures favoring larger brain size often appear more pronounced in non-reproductive contexts. For instance, habitat complexity may favor larger-brained juveniles by enhancing predator avoidance strategies as structural features of complex environments provide better opportunities for evading predators (Johnson, 2007). In contrast, open environments may exert different sexual selection pressures, particularly by providing greater opportunities for mate assessment. The specialized surface dwelling halfbeak fish we examined may be particularly adapted to mating conditions in relatively open environments, as adult halfbeaks live in shoals that rarely venture deeper than a few centimeters below the surface of the water (Devigili et al., 2021). Under these conditions, both males and females may need to constantly process and compare information about mating opportunities. As a result, cognitive processes relevant for fitness become more important in open environments where the potential to assess mates increases. Supporting this idea, the only observed link between female gravid spot size, which males use during mate choice to assess female reproductive receptivity, and female fitness was found in open arenas. Additionally, bold behaviors have been shown to provide individuals with preferential access to mates in open settings (Myhre et al., 2013), and brain size has been positively correlated with proactive behaviors in other fishes (e.g., guppies Kotrschal et al., 2014*b*). Open environments are expected to facilitate shoal formation and maintenance which in turn, may increase the rate of social behaviors, especially agonistic interactions in this species. By facilitating proactive behaviors, such as boldness, a larger brain may enable individuals to outcompete rivals and court more mates in open settings. Conversely, in structurally complex habitats, where social interaction may be more scarce due to physical or visual constraints, boldness may confer fewer reproductive advantages. Open environments, while facilitating mate assessment, may also represent higher perceived predation risk in natural settings for surface-dwelling species like halfbeaks. Larger brain size has been linked to improved survival under predation threat in other fish species (e.g., Kotrschal et al., 2015), raising the possibility that stronger selection on brain size in open environments could also reflect behavioral strategies shaped by natural selection, even in the absence of direct predation risk. Our findings highlight the need to further investigate how habitat complexity shapes selection on brain morphology, particularly by disentangling the relative fitness benefits of neural investment in reproductive versus non-reproductive contexts and under ecologically realistic conditions that include predation risk.

Interestingly, we found little evidence of selection on male ornamentation or body size in either environment. The only evidence of selection on male ornaments was in complex environments, where we detected weak non-linear stabilizing selection on red coloration, suggesting that males with intermediate levels of red coloration are favored over those with very low or very high levels. Previous studies on halfbeaks using dichotomous mate choice tests have shown that female preference for red coloration is variable and context-dependent. For example, female preference differs between virgin and mated females (Reuland et al., 2020) and depends on the absolute magnitude of color difference between males (McNeil et al., 2021). Such variation in female preference may explain the pattern of stabilizing selection on red coloration detected in our study. Alternatively, intermediately colored males may strike a balance between being noticeable enough for mate attraction while not being too conspicuous to rival males. Thus, males with intermediate coloration might have an advantage in terms of both avoiding direct male-male competition (which may be more intense for showy males in complex environment, see below) and being attractive enough to females compared to the least red males. In line with this argument, in sticklebacks (*Gasterosteus aculeatus*) the amount of red coloration on a male directly affects a male’s aggression success (Lackey and Boughman, 2013), highlighting that colorful males dominated the others through physical contests. Moreover, some studies have shown that male-male competition may be higher in complex environments with small enclosed groups than in open, high density, unstructured environments (e.g., Lukasik et al., 2006). In such cases, highly competitive males in complex environments may pay fitness costs of higher aggressiveness, if the time spent being aggressive does not ensure mate monopolization. However, our study cannot properly assess whether stabilizing selection on red coloration results from female mate choice or male competition. Furthermore, the lack of selection on male body and beak size in both arena types prevents drawing conclusions about differences in the intensity of male competition between open and complex environments. We recommend that future research directly explores male competition across different habitat types, as this would provide valuable insights into how environmental factors influence male-male competition and how success in this competition relates to morphological traits.

In conclusion, our study demonstrates that habitat complexity is a key factor interacting with brain size to shape fitness in halfbeaks. The positive linear selection gradients on brain size detected in female and male halfbeaks suggest that sexually selected processes can drive brain evolution. The direction of selection arising from naturally and sexually selected processes may thus align, resulting in increased evolutionary pressure on brain size. However, predicting the environmental conditions where selection will favor increased brain size remains challenging as we found little evidence of selection on brain size in complex environments despite clear theoretical predictions. More broadly, our results (1) reinforce recent arguments emphasizing the need to better integrate cognitive processes into studies of mate choice and sexual selection (Ryan, 2021) and (2) emphasize the importance of quantifying sexual selection metrics in both sexes to fully understand how pre-copulatory sexual selection operates. Future studies should also consider how post-copulatory processes may interact with cognitive traits and habitat complexity to shape fitness outcomes. By considering selection gradients and other quantitative metrics for both males and females across a range of ecologically relevant conditions, we can gain a more nuanced perspective of how selective pressures act across different traits and contexts within a system.

## Authors’ contributions

L.D.: statistical analysis, writing—original draft, writing—review and editing; A.D.: conceptualisation, methodology, data curation, writing—review and editing; R.M.: methodology, data curation, editing; N. K. and D.W.: conceptualisation, editing; J.F.: conceptualisation, funding acquisition, methodology, supervision, writing—original draft, writing—review and editing. All authors gave final approval for publication and agreed to be held accountable for the work performed therein.

## Acknowledgments

We thank all current and past members of Fitzpatrick Lab, especially Mirjam Amcoff, for their great support in fish feeding and monitoring. This work was funded by a Swedish Research Council Grant (2021-04615) to J.L.F. and a Carl Tryggers Foundation postdoctoral stipend (CTS 23:2794) to L.D.

## Data accessibility

Data and scripts will be available on Dryad.

## Appendix A Supplemental Information

**Table S1:**
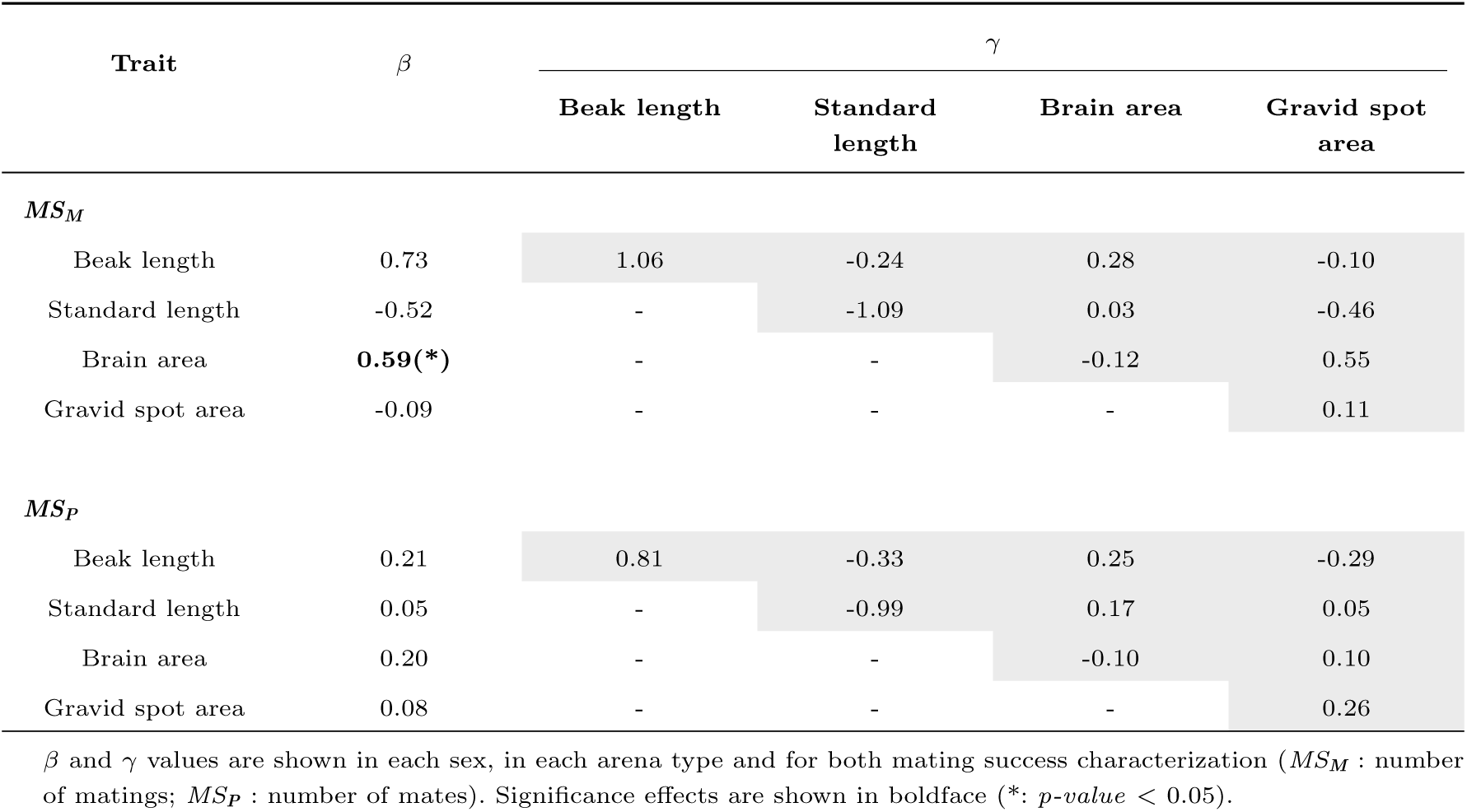
Linear *β* and non-linear (quadratic and correlational, *γ*) selection gradients in females in open arenas.

**Table S2:**
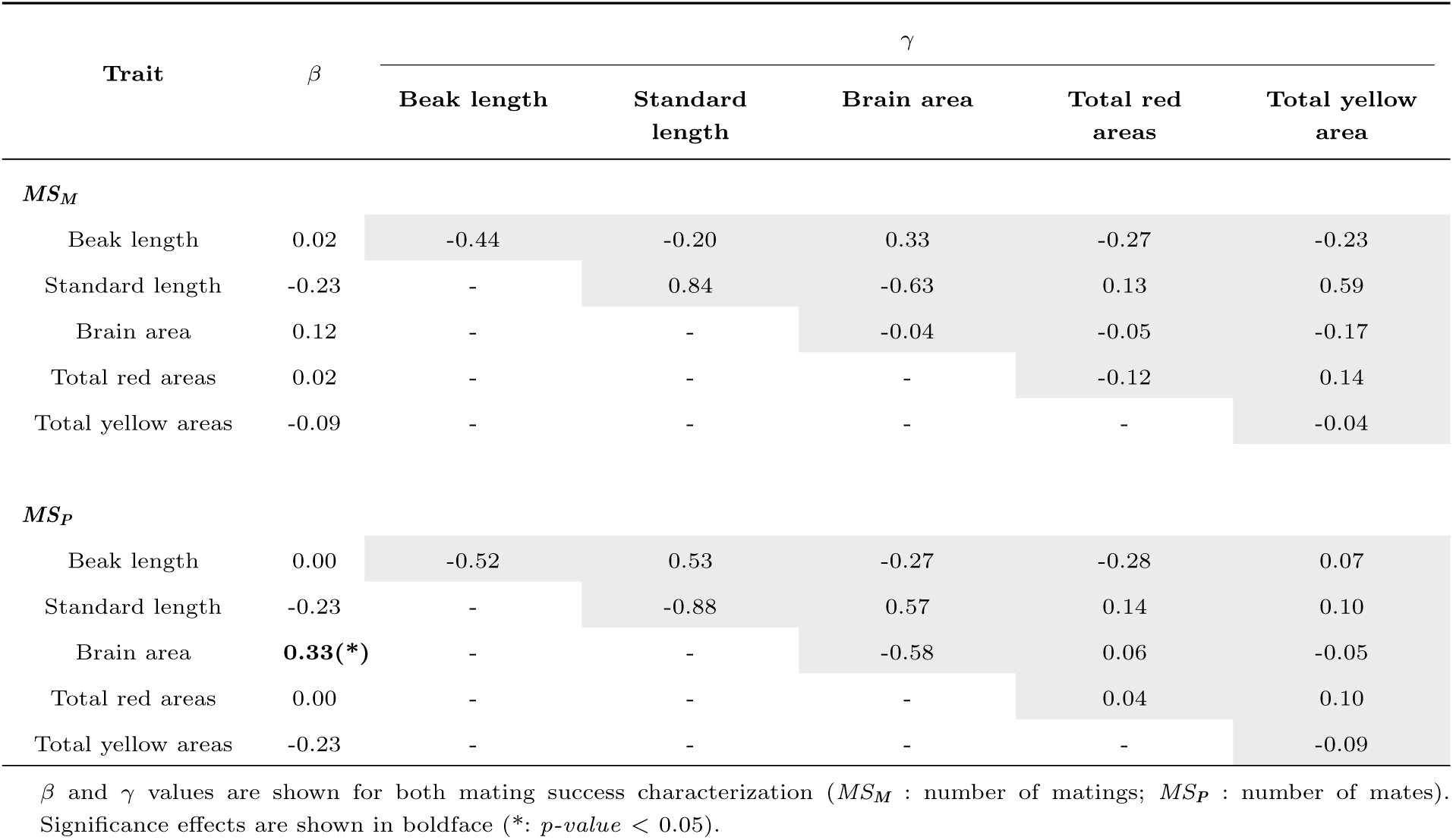
Linear *β* and non-linear (quadratic and correlational, *γ*) selection gradients in males in open arenas.

**Table S3:**
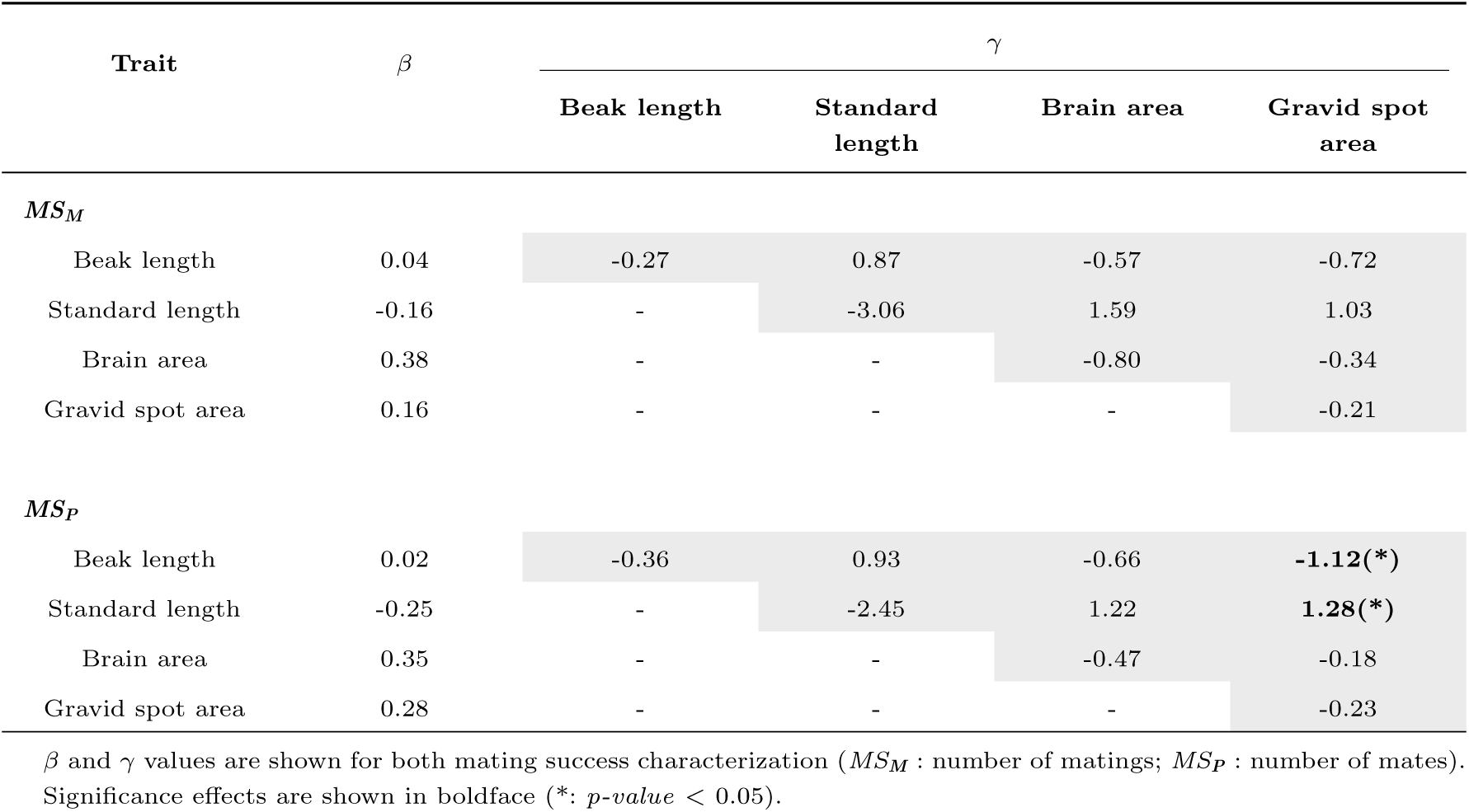
Linear *β* and non-linear (quadratic and correlational, *γ*) selection gradients in females in complex arenas.

**Table S4:**
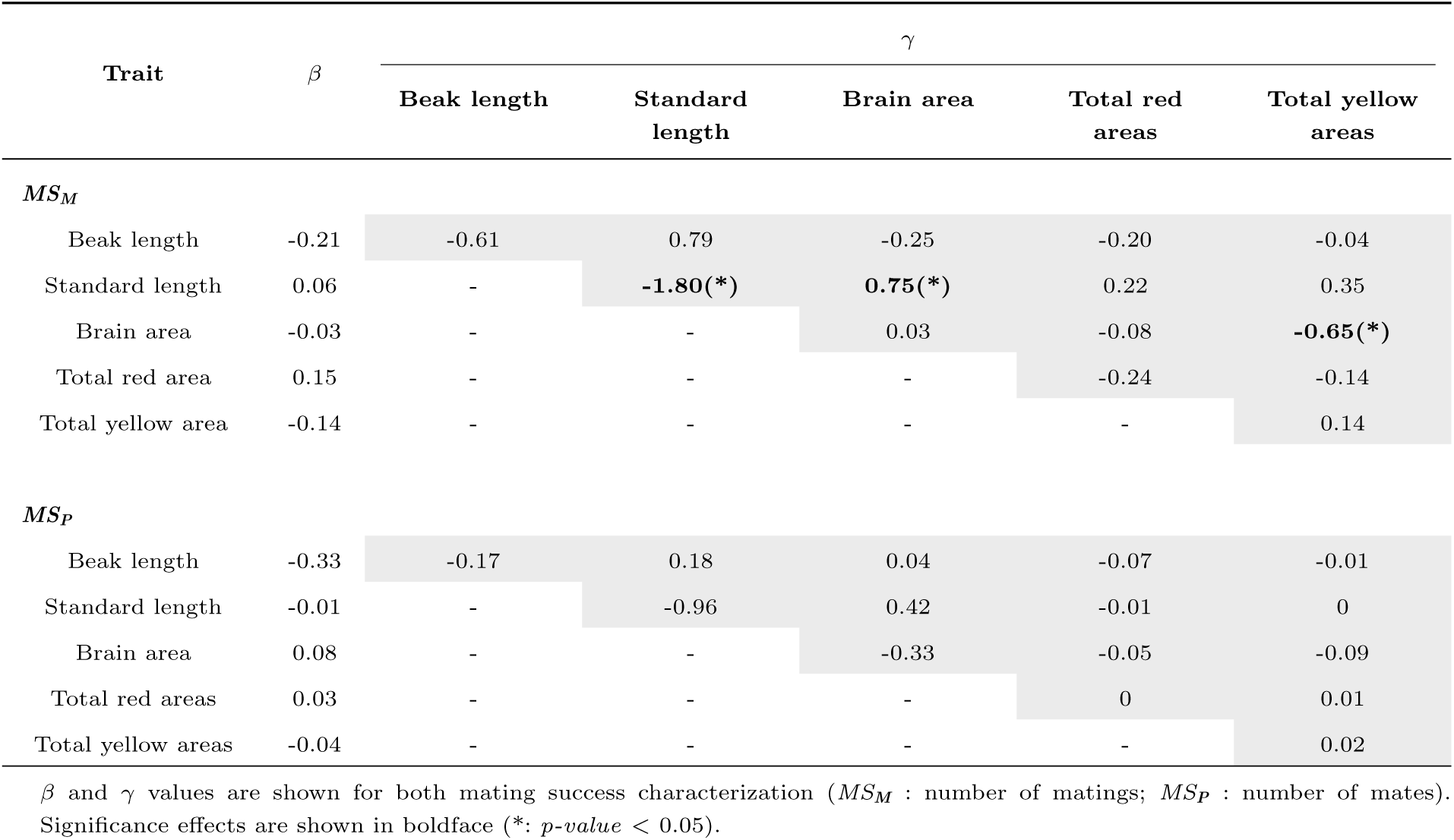
Linear *β* and non-linear (quadratic and correlational, *γ*) selection gradients in males in complex arenas.

**Table S5:**
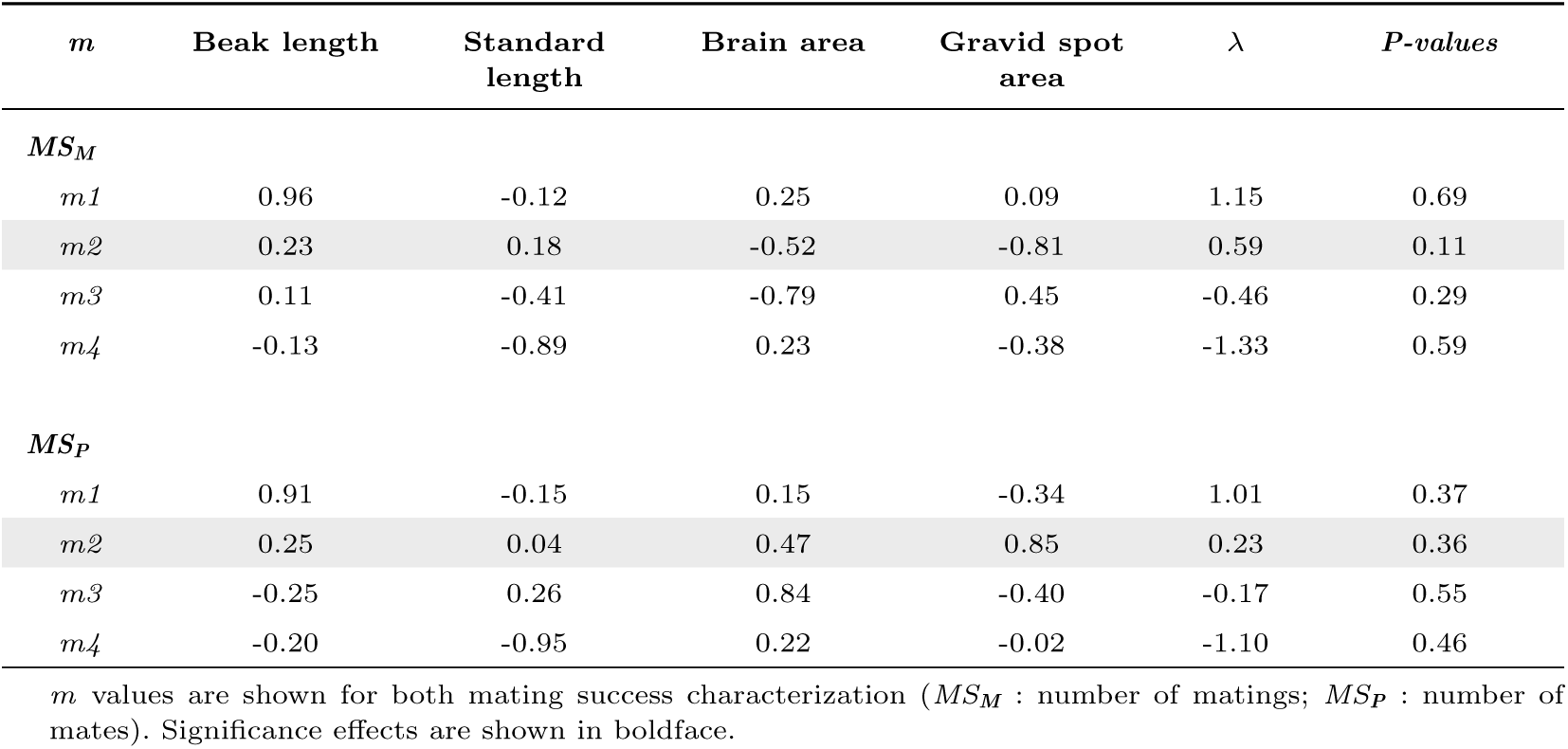
The matrix of eigenvectors (*m*) and estimates of non-linear selection on the axes (eigenvalues, *λ*) described by the eigenvectors from the canonical analysis of gamma matrix *γ* in females in open arenas.

**Table S6:**
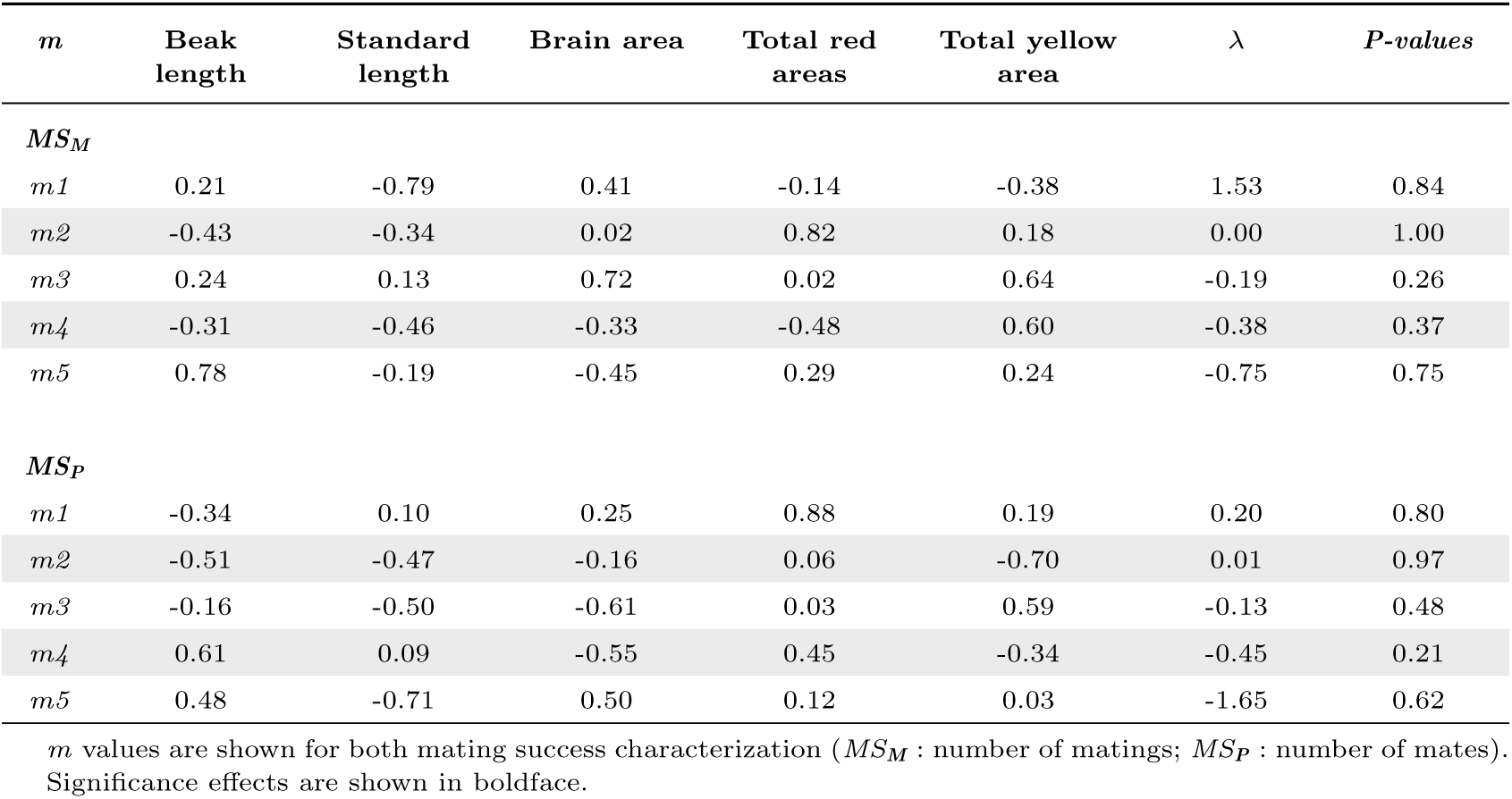
The matrix of eigenvectors (*m*) and estimates of non-linear selection on the axes (eigenvalues, *λ*) described by the eigenvectors from the canonical analysis of gamma matrix *γ* in males in open arenas.

**Table S7:**
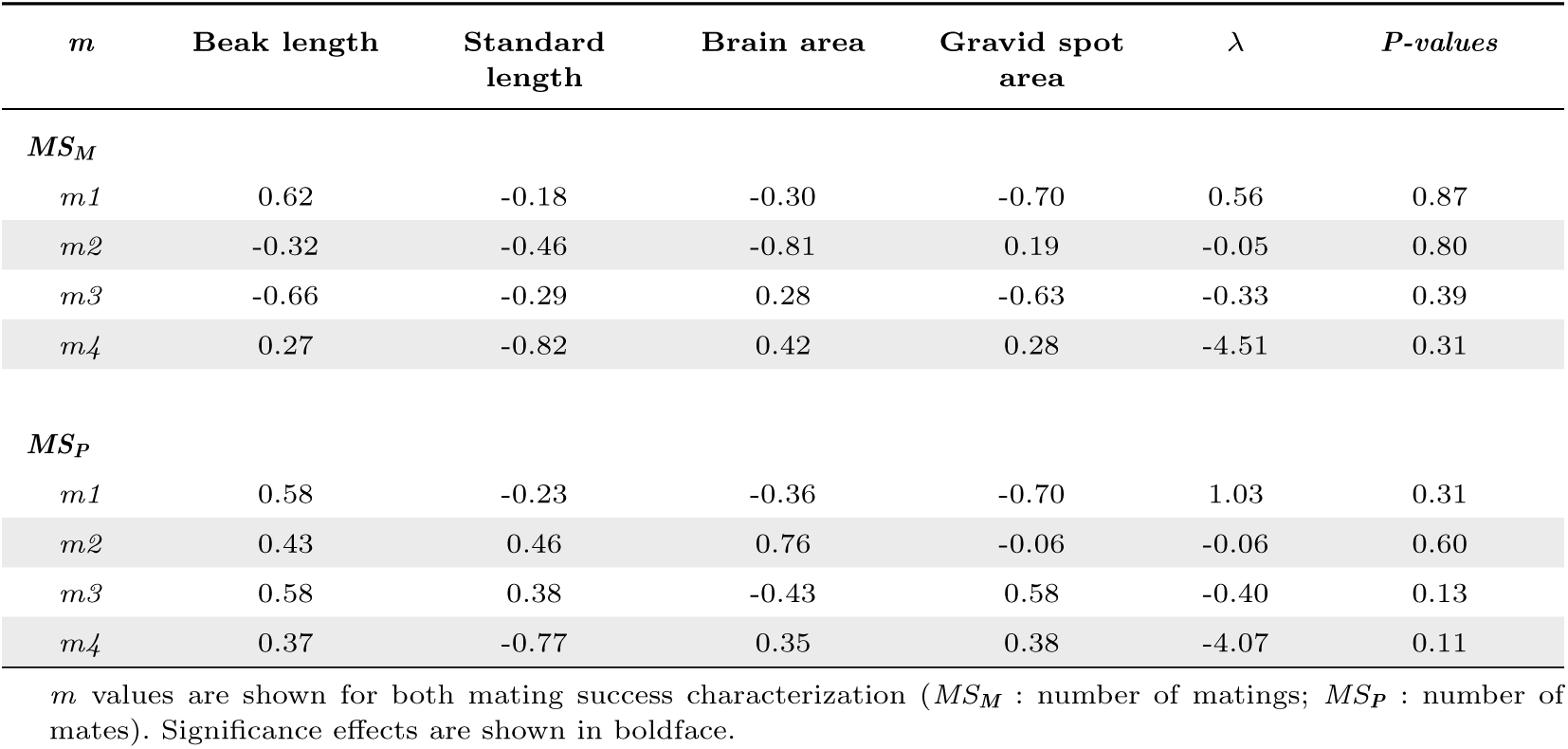
The matrix of eigenvectors (*m*) and estimates of non-linear selection on the axes (eigenvalues, *λ*) described by the eigenvectors from the canonical analysis of gamma matrix *γ* in females in complex arenas.

**Table S8:**
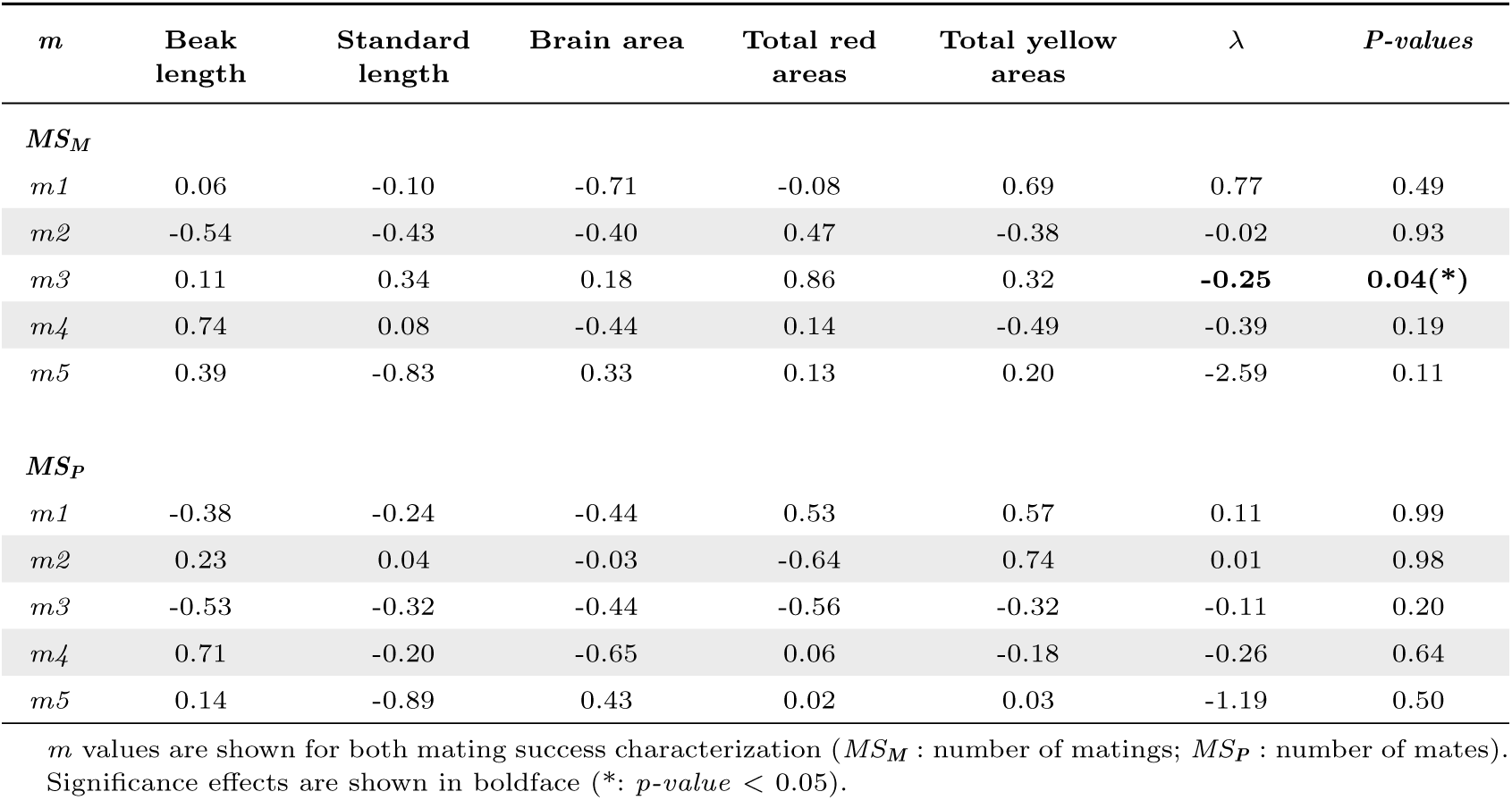
The matrix of eigenvectors (*m*) and estimates of non-linear selection on the axes (eigenvalues, *λ*) described by the eigenvectors from the canonical analysis of gamma matrix *γ* in males in complex arenas.

**Table S9:**
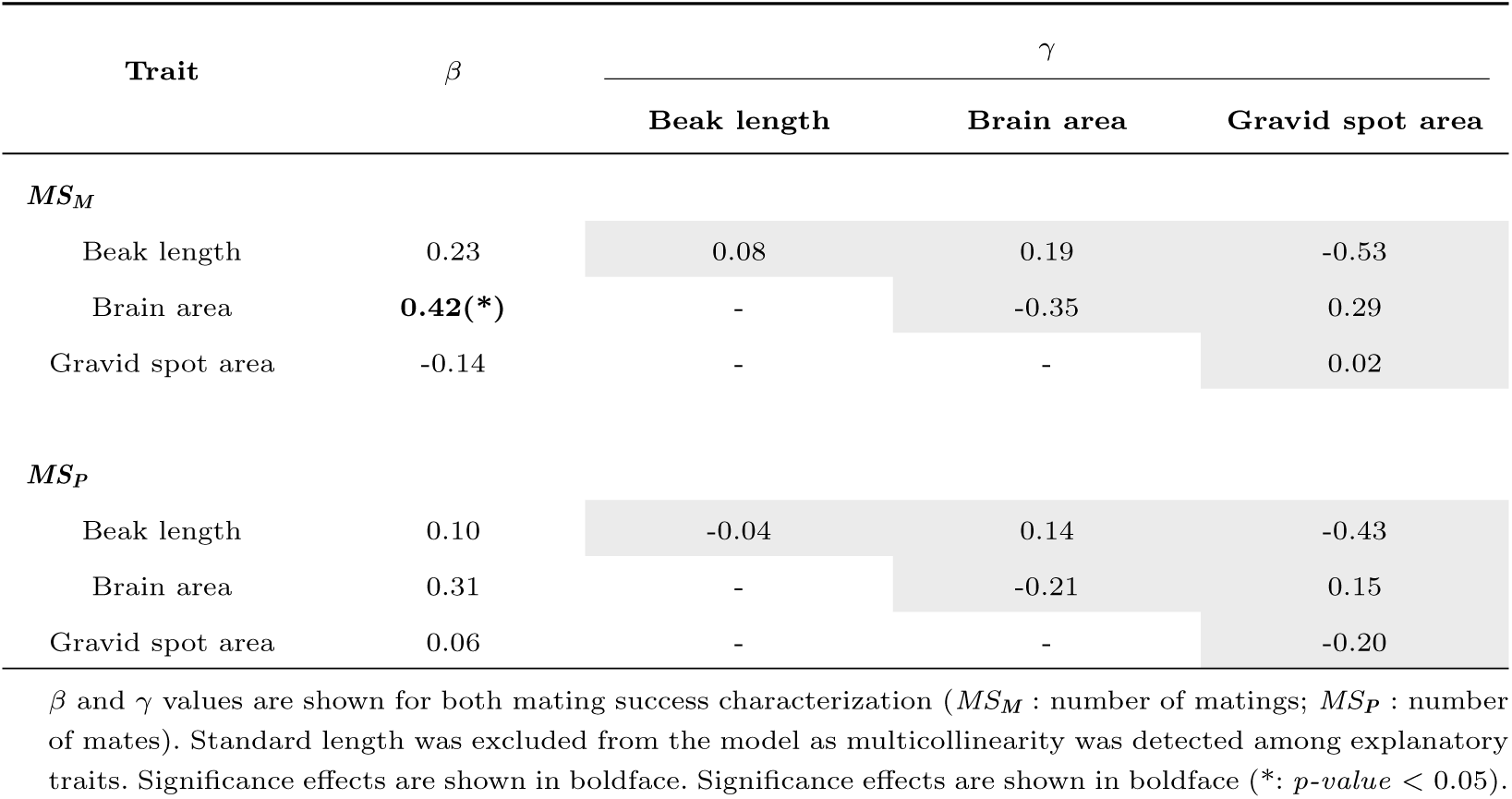
Linear *β* and non-linear (quadratic and correlational, *γ*) selection gradients in females in open arenas.

**Table S10:**
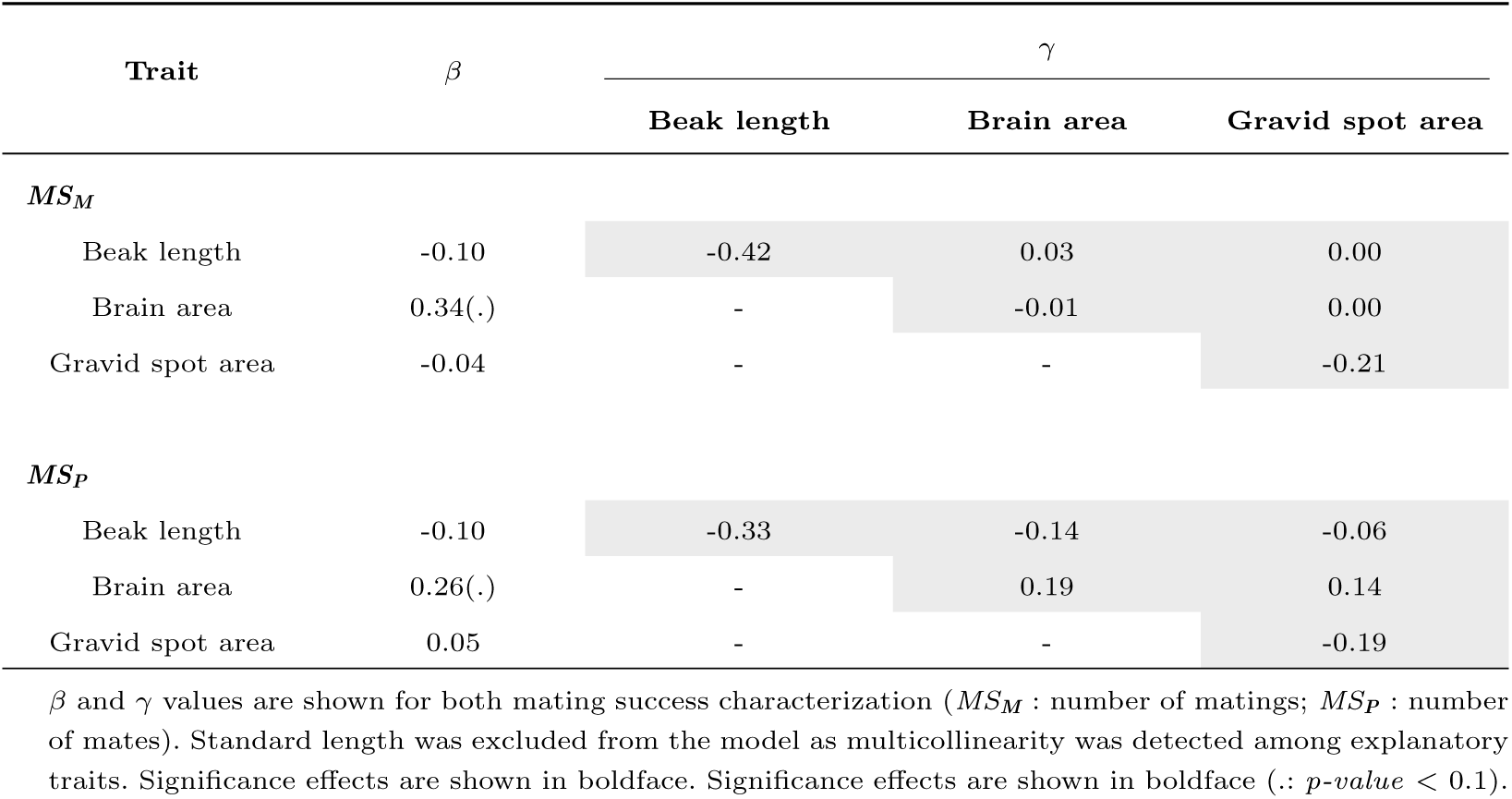
Linear *β* and non-linear (quadratic and correlational, *γ*) selection gradients in females in complex arenas.

**Table S11:**
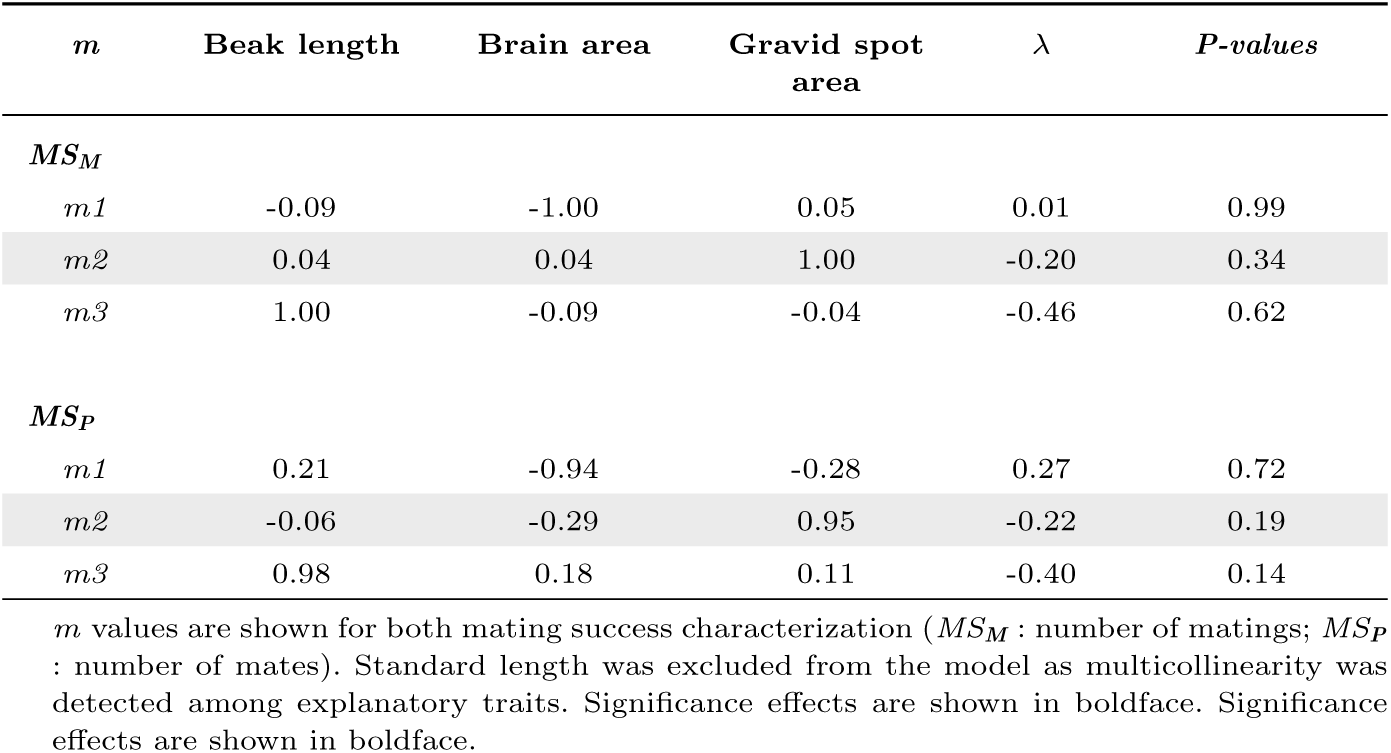
The matrix of eigenvectors (*m*) and estimates of non-linear selection on the axes (eigenvalues, *λ*) described by the eigenvectors from the canonical analysis of gamma matrix *γ* in females in complex arenas.

**Table S12:**
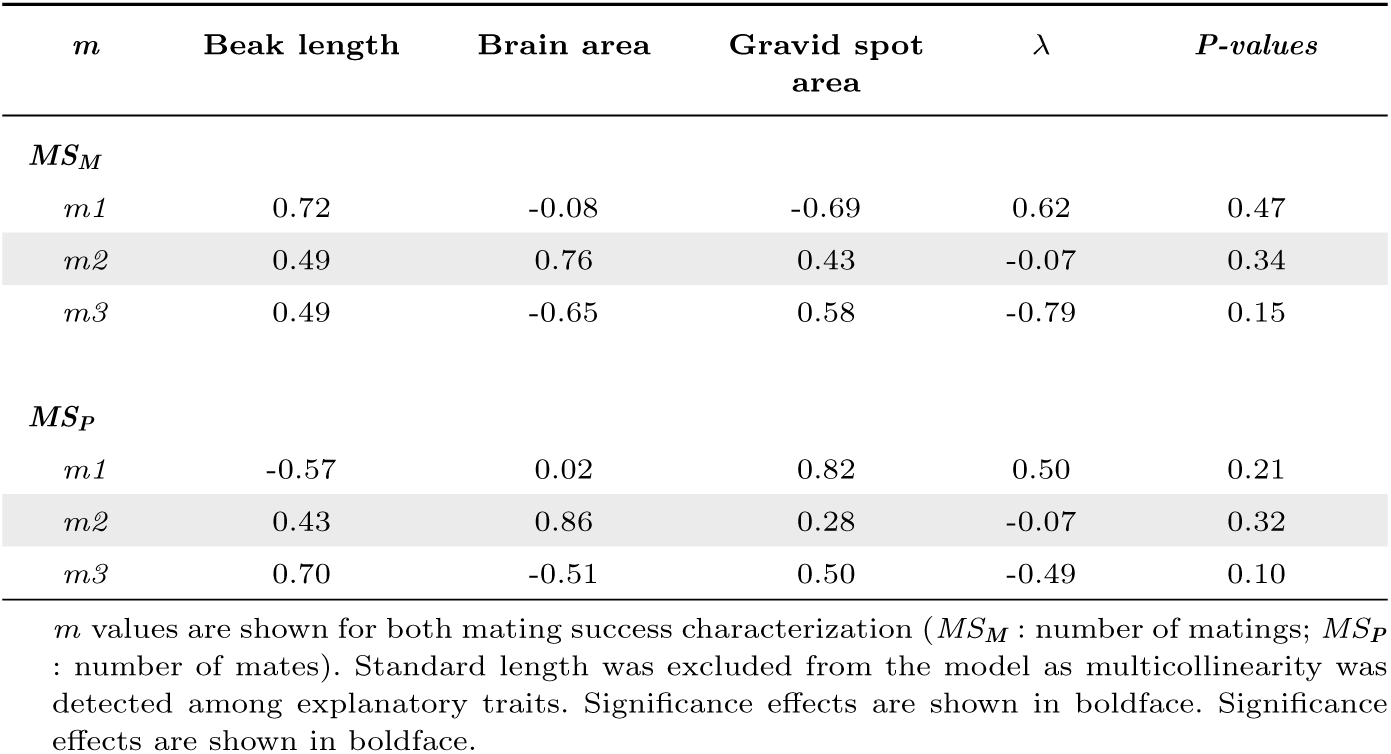
The matrix of eigenvectors (*m*) and estimates of non-linear selection on the axes (eigenvalues, *λ*) described by the eigenvectors from the canonical analysis of gamma matrix *γ* in females in open arenas.

**Fig. S1:**
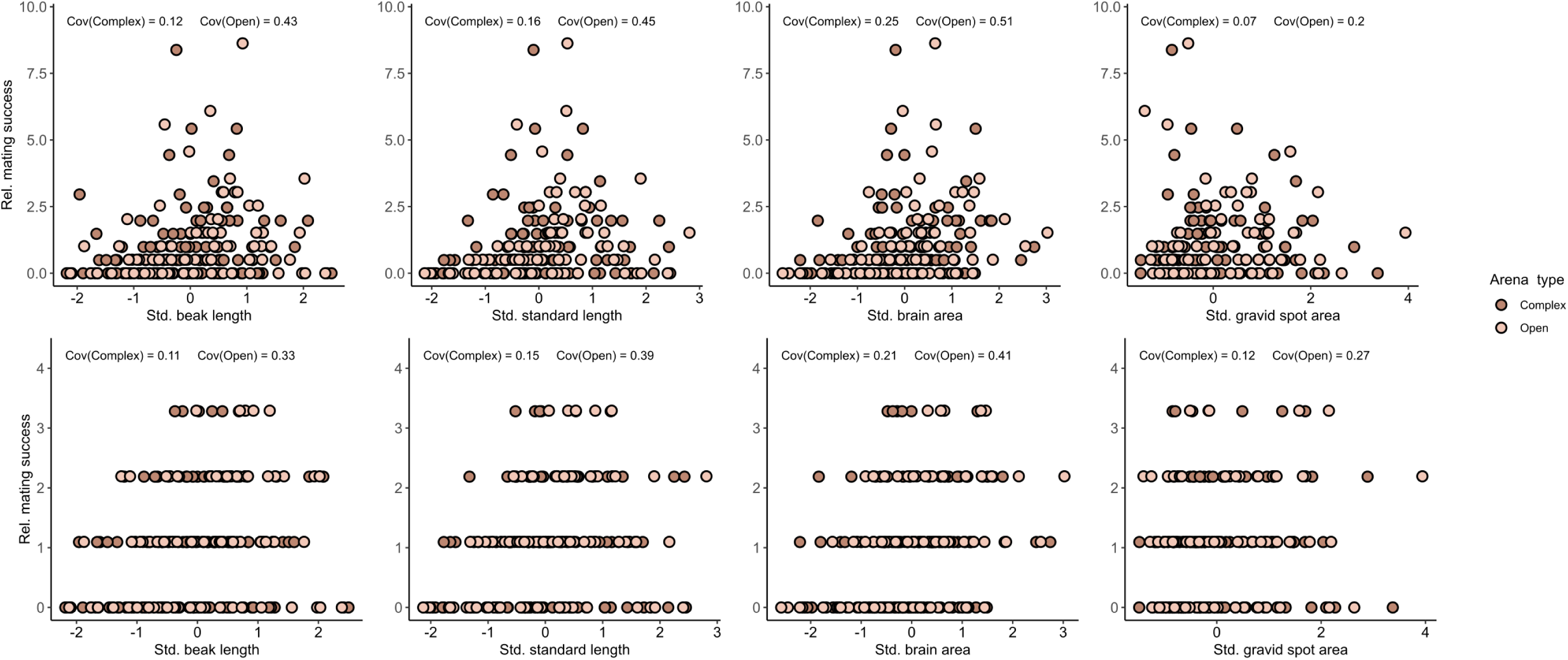
Relationship between standardized traits and relative mating success success for females in open (light pink) and (dark pink) arenas. Covariance values obtained in each arena type are indicated above each plot.

**Fig. S2:**
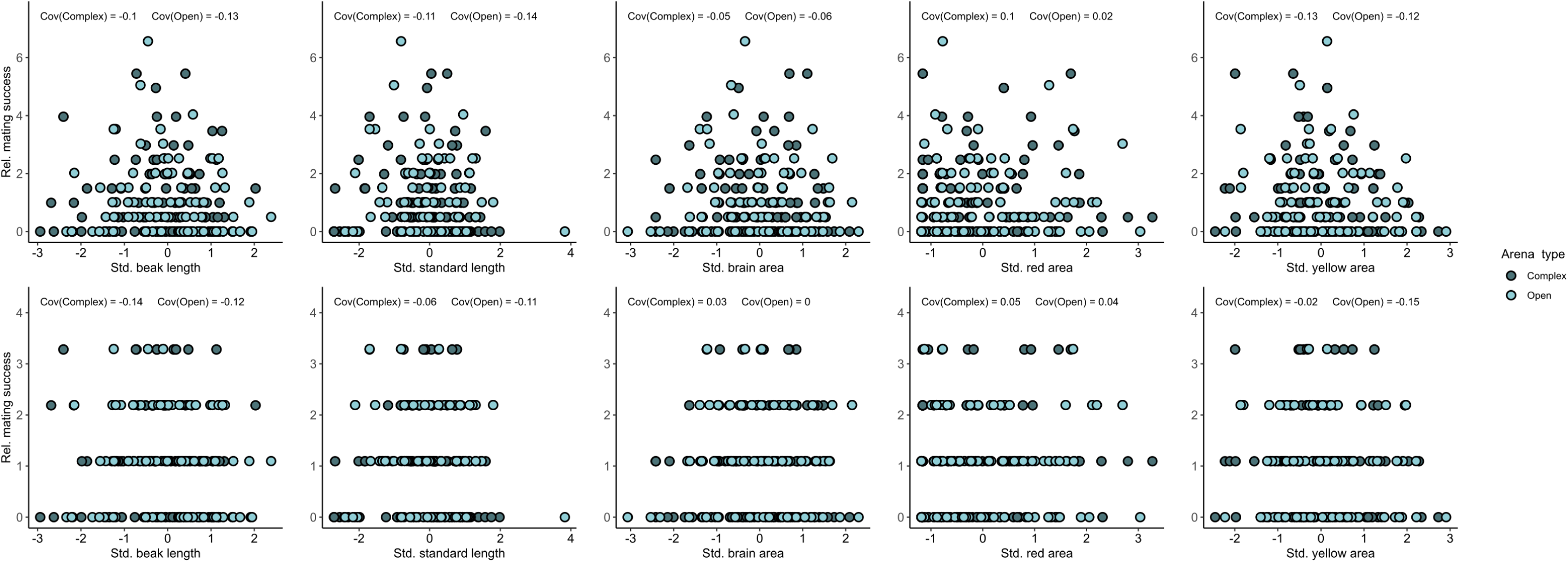
Relationship between standardized traits and relative mating success success for males in open (light blue) and (dark blue) arenas. Covariance values obtained in each arena type are indicated above each plot.

**Table S13:**
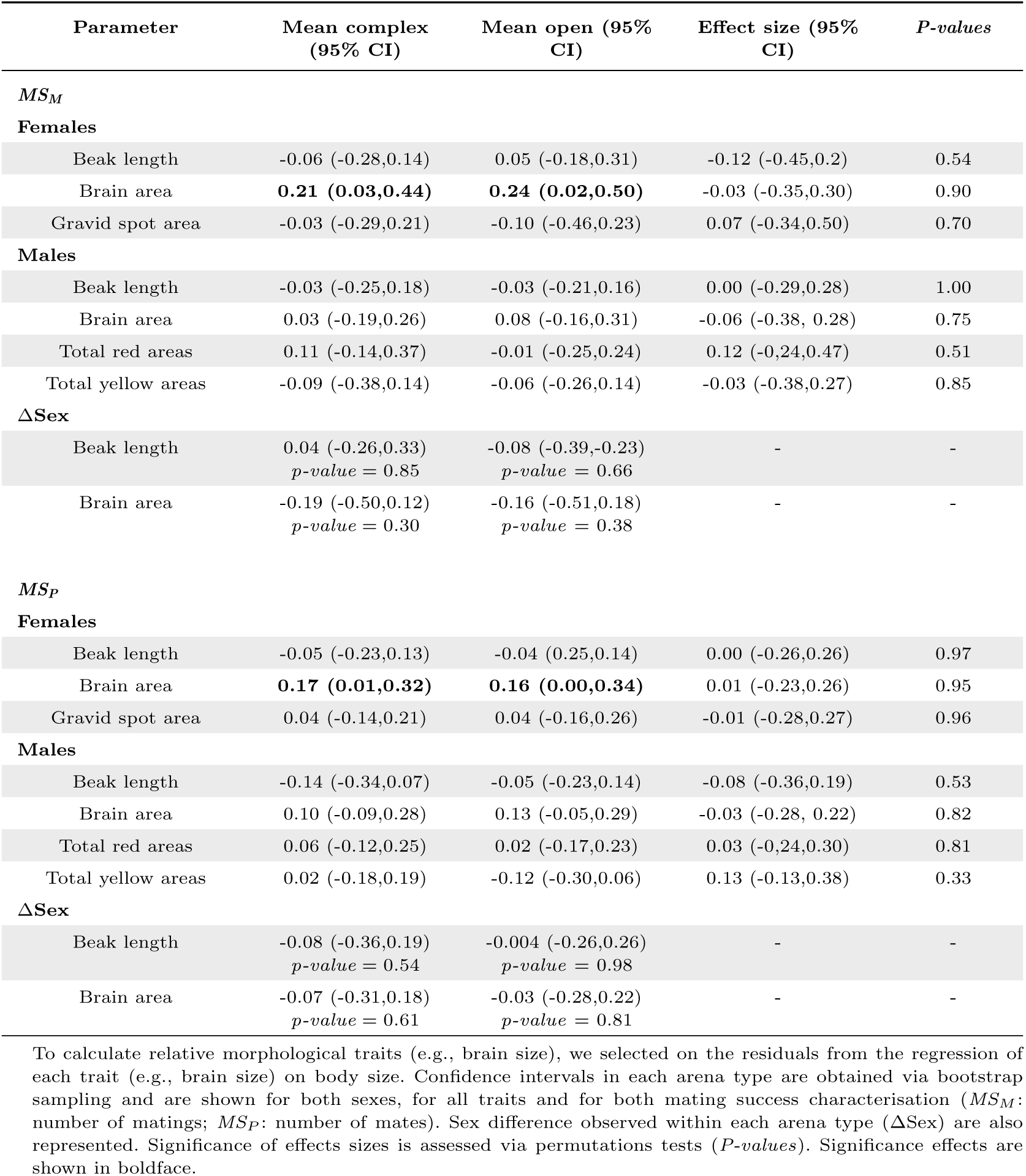
Influence of arena type on mating differentials (*m’*) on relative morphological traits.

