## Supplementary Material for "Habitat complexity alters the strength of sexual selection on brain size in a livebearing fish"

**Table S1**: Linear $\beta$ and non-linear (quadratic and correlational, $\gamma$) selection gradients in **females** in **open arenas**

| **Trait** | $\beta$ | $\gamma$ | | | |
|---|---|---|---|---|---|
| | | **Beak length** | **Standard length** | **Brain area** | **Gravid spot area** |
| $\boldsymbol{MS_M}$ | | | | | |
| Beak length | 0.73 | 1.06 | -0.24 | 0.28 | -0.10 |
| Standard length | -0.52 | - | -1.09 | 0.03 | -0.46 |
| Brain area | **0.59(*)** | - | - | -0.12 | 0.55 |
| Gravid spot area | -0.09 | - | - | - | 0.11 |
| $\boldsymbol{MS_P}$ | | | | | |
| Beak length | 0.21 | 0.81 | -0.33 | 0.25 | -0.29 |
| Standard length | 0.05 | - | -0.99 | 0.17 | 0.05 |
| Brain area | 0.20 | - | - | -0.10 | 0.10 |
| Gravid spot area | 0.08 | - | - | - | 0.26 |

$\beta$ and $\gamma$ values are shown in each sex, in each arena type and for both mating success characterization ($MS_M$ : number of matings; $MS_P$ : number of mates). Significance effects are shown in boldface (*: *p-value* $< 0.05$).

**Table S2**: Linear $\beta$ and non-linear (quadratic and correlational, $\gamma$) selection gradients in **males** in **open arenas**

| **Trait** | $\beta$ | $\gamma$ | | | | |
|---|---|---|---|---|---|---|
| | | **Beak length** | **Standard length** | **Brain area** | **Total red areas** | **Total yellow area** |
| ***$MS_M$*** | | | | | | |
| Beak length | 0.02 | -0.44 | -0.20 | 0.33 | -0.27 | -0.23 |
| Standard length | -0.23 | - | 0.84 | -0.63 | 0.13 | 0.59 |
| Brain area | 0.12 | - | - | -0.04 | -0.05 | -0.17 |
| Total red areas | 0.02 | - | - | - | -0.12 | 0.14 |
| Total yellow areas | -0.09 | - | - | - | - | -0.04 |
| ***$MS_P$*** | | | | | | |
| Beak length | 0.00 | -0.52 | 0.53 | -0.27 | -0.28 | 0.07 |
| Standard length | -0.23 | - | -0.88 | 0.57 | 0.14 | 0.10 |
| Brain area | **0.33(*)** | - | - | -0.58 | 0.06 | -0.05 |
| Total red areas | 0.00 | - | - | - | 0.04 | 0.10 |
| Total yellow areas | -0.23 | - | - | - | - | -0.09 |

$\beta$ and $\gamma$ values are shown for both mating success characterization ($MS_M$ : number of matings; $MS_P$ : number of mates). Significance effects are shown in boldface (*: *p-value* $< 0.05$).

**Table S3**: Linear $\beta$ and non-linear (quadratic and correlational, $\gamma$) selection gradients in **females** in **complex arenas**

| **Trait** | $\beta$ | $\gamma$ | | | |
|---|---|---|---|---|---|
| | | **Beak length** | **Standard length** | **Brain area** | **Gravid spot area** |
| ***$MS_M$*** | | | | | |
| Beak length | 0.04 | -0.27 | 0.87 | -0.57 | -0.72 |
| Standard length | -0.16 | - | -3.06 | 1.59 | 1.03 |
| Brain area | 0.38 | - | - | -0.80 | -0.34 |
| Gravid spot area | 0.16 | - | - | - | -0.21 |
| ***$MS_P$*** | | | | | |
| Beak length | 0.02 | -0.36 | 0.93 | -0.66 | **-1.12(*)** |
| Standard length | -0.25 | - | -2.45 | 1.22 | **1.28(*)** |
| Brain area | 0.35 | - | - | -0.47 | -0.18 |
| Gravid spot area | 0.28 | - | - | - | -0.23 |

$\beta$ and $\gamma$ values are shown for both mating success characterization ($MS_M$ : number of matings; $MS_P$ : number of mates). Significance effects are shown in boldface (*: *p-value* $< 0.05$).

**Table S4**: Linear $\beta$ and non-linear (quadratic and correlational, $\gamma$) selection gradients in **males** in **complex arenas**

| **Trait** | $\beta$ | $\gamma$ | | | | |
|---|---|---|---|---|---|---|
| | | **Beak length** | **Standard length** | **Brain area** | **Total red areas** | **Total yellow areas** |
| ***$MS_M$*** | | | | | | |
| Beak length | -0.21 | -0.61 | 0.79 | -0.25 | -0.20 | -0.04 |
| Standard length | 0.06 | - | **-1.80(*)** | **0.75(*)** | 0.22 | 0.35 |
| Brain area | -0.03 | - | - | 0.03 | -0.08 | **-0.65(*)** |
| Total red area | 0.15 | - | - | - | -0.24 | -0.14 |
| Total yellow area | -0.14 | - | - | - | - | 0.14 |
| ***$MS_P$*** | | | | | | |
| Beak length | -0.33 | -0.17 | 0.18 | 0.04 | -0.07 | -0.01 |
| Standard length | -0.01 | - | -0.96 | 0.42 | -0.01 | 0 |
| Brain area | 0.08 | - | - | -0.33 | -0.05 | -0.09 |
| Total red areas | 0.03 | - | - | - | 0 | 0.01 |
| Total yellow areas | -0.04 | - | - | - | - | 0.02 |

$\beta$ and $\gamma$ values are shown for both mating success characterization ($MS_M$ : number of matings; $MS_P$ : number of mates). Significance effects are shown in boldface (*: *p-value* $< 0.05$).

**Table S5**: The matrix of eigenvectors ($m$) and estimates of non-linear selection on the axes (eigenvalues, $\lambda$) described by the eigenvectors from the canonical analysis of gamma matrix $\gamma$ in **females** in **open arenas**

| $m$ | **Beak length** | **Standard length** | **Brain area** | **Gravid spot area** | $\lambda$ | ***P-values*** |
|---|---|---|---|---|---|---|
| ***$MS_M$*** | | | | | | |
| *m1* | 0.96 | -0.12 | 0.25 | 0.09 | 1.15 | 0.69 |
| *m2* | 0.23 | 0.18 | -0.52 | -0.81 | 0.59 | 0.11 |
| *m3* | 0.11 | -0.41 | -0.79 | 0.45 | -0.46 | 0.29 |
| *m4* | -0.13 | -0.89 | 0.23 | -0.38 | -1.33 | 0.59 |
| ***$MS_P$*** | | | | | | |
| *m1* | 0.91 | -0.15 | 0.15 | -0.34 | 1.01 | 0.37 |
| *m2* | 0.25 | 0.04 | 0.47 | 0.85 | 0.23 | 0.36 |
| *m3* | -0.25 | 0.26 | 0.84 | -0.40 | -0.17 | 0.55 |
| *m4* | -0.20 | -0.95 | 0.22 | -0.02 | -1.10 | 0.46 |

$m$ values are shown for both mating success characterization ($MS_M$ : number of matings; $MS_P$ : number of mates). Significance effects are shown in boldface.

**Table S6**: The matrix of eigenvectors ($m$) and estimates of non-linear selection on the axes (eigenvalues, $\lambda$) described by the eigenvectors from the canonical analysis of gamma matrix $\gamma$ in **males** in **open arenas**

| ***m*** | **Beak length** | **Standard length** | **Brain area** | **Total red areas** | **Total yellow area** | $\lambda$ | ***P-values*** |
|---|---|---|---|---|---|---|---|
| $MS_M$ | | | | | | | |
| *m1* | 0.21 | -0.79 | 0.41 | -0.14 | -0.38 | 1.53 | 0.84 |
| *m2* | -0.43 | -0.34 | 0.02 | 0.82 | 0.18 | 0.00 | 1.00 |
| *m3* | 0.24 | 0.13 | 0.72 | 0.02 | 0.64 | -0.19 | 0.26 |
| *m4* | -0.31 | -0.46 | -0.33 | -0.48 | 0.60 | -0.38 | 0.37 |
| *m5* | 0.78 | -0.19 | -0.45 | 0.29 | 0.24 | -0.75 | 0.75 |
| $MS_P$ | | | | | | | |
| *m1* | -0.34 | 0.10 | 0.25 | 0.88 | 0.19 | 0.20 | 0.80 |
| *m2* | -0.51 | -0.47 | -0.16 | 0.06 | -0.70 | 0.01 | 0.97 |
| *m3* | -0.16 | -0.50 | -0.61 | 0.03 | 0.59 | -0.13 | 0.48 |
| *m4* | 0.61 | 0.09 | -0.55 | 0.45 | -0.34 | -0.45 | 0.21 |
| *m5* | 0.48 | -0.71 | 0.50 | 0.12 | 0.03 | -1.65 | 0.62 |

$m$ values are shown for both mating success characterization ($MS_M$ : number of matings; $MS_P$ : number of mates). Significance effects are shown in boldface.

**Table S7**: The matrix of eigenvectors ($m$) and estimates of non-linear selection on the axes (eigenvalues, $\lambda$) described by the eigenvectors from the canonical analysis of gamma matrix $\gamma$ in **females** in **complex arenas**

| ***m*** | **Beak length** | **Standard length** | **Brain area** | **Gravid spot area** | $\lambda$ | ***P-values*** |
|---|---|---|---|---|---|---|
| $MS_M$ | | | | | | |
| *m1* | 0.62 | -0.18 | -0.30 | -0.70 | 0.56 | 0.87 |
| *m2* | -0.32 | -0.46 | -0.81 | 0.19 | -0.05 | 0.80 |
| *m3* | -0.66 | -0.29 | 0.28 | -0.63 | -0.33 | 0.39 |
| *m4* | 0.27 | -0.82 | 0.42 | 0.28 | -4.51 | 0.31 |
| $MS_P$ | | | | | | |
| *m1* | 0.58 | -0.23 | -0.36 | -0.70 | 1.03 | 0.31 |
| *m2* | 0.43 | 0.46 | 0.76 | -0.06 | -0.06 | 0.60 |
| *m3* | 0.58 | 0.38 | -0.43 | 0.58 | -0.40 | 0.13 |
| *m4* | 0.37 | -0.77 | 0.35 | 0.38 | -4.07 | 0.11 |

$m$ values are shown for both mating success characterization ($MS_M$ : number of matings; $MS_P$ : number of mates). Significance effects are shown in boldface.

**Table S8**: The matrix of eigenvectors ($m$) and estimates of non-linear selection on the axes (eigenvalues, $\lambda$) described by the eigenvectors from the canonical analysis of gamma matrix $\gamma$ in **males** in **complex arenas**

| $m$ | **Beak length** | **Standard length** | **Brain area** | **Total red areas** | **Total yellow areas** | $\lambda$ | ***P-values*** |
|---|---|---|---|---|---|---|---|
| $MS_M$ | | | | | | | |
| *m1* | 0.06 | -0.10 | -0.71 | -0.08 | 0.69 | 0.77 | 0.49 |
| *m2* | -0.54 | -0.43 | -0.40 | 0.47 | -0.38 | -0.02 | 0.93 |
| *m3* | 0.11 | 0.34 | 0.18 | 0.86 | 0.32 | **-0.25** | **0.04(*)** |
| *m4* | 0.74 | 0.08 | -0.44 | 0.14 | -0.49 | -0.39 | 0.19 |
| *m5* | 0.39 | -0.83 | 0.33 | 0.13 | 0.20 | -2.59 | 0.11 |
| $MS_P$ | | | | | | | |
| *m1* | -0.38 | -0.24 | -0.44 | 0.53 | 0.57 | 0.11 | 0.99 |
| *m2* | 0.23 | 0.04 | -0.03 | -0.64 | 0.74 | 0.01 | 0.98 |
| *m3* | -0.53 | -0.32 | -0.44 | -0.56 | -0.32 | -0.11 | 0.20 |
| *m4* | 0.71 | -0.20 | -0.65 | 0.06 | -0.18 | -0.26 | 0.64 |
| *m5* | 0.14 | -0.89 | 0.43 | 0.02 | 0.03 | -1.19 | 0.50 |

$m$ values are shown for both mating success characterization ($MS_M$ : number of matings; $MS_P$ : number of mates). Significance effects are shown in boldface (*: *p-value* $< 0.05$).

**Table S9**: Linear $\beta$ and non-linear (quadratic and correlational, $\gamma$) selection gradients in **females** in **open arenas**

| **Trait** | $\beta$ | $\gamma$ | | |
|---|---|---|---|---|
| | | **Beak length** | **Brain area** | **Gravid spot area** |
| $MS_M$ | | | | |
| Beak length | 0.23 | 0.08 | 0.19 | -0.53 |
| Brain area | **0.42(*)** | - | -0.35 | 0.29 |
| Gravid spot area | -0.14 | - | - | 0.02 |
| $MS_P$ | | | | |
| Beak length | 0.10 | -0.04 | 0.14 | -0.43 |
| Brain area | 0.31 | - | -0.21 | 0.15 |
| Gravid spot area | 0.06 | - | - | -0.20 |

$\beta$ and $\gamma$ values are shown for both mating success characterization ($MS_M$ : number of matings; $MS_P$ : number of mates). Standard length was excluded from the model as multicollinearity was detected among explanatory traits. Significance effects are shown in boldface. Significance effects are shown in boldface (*: *p-value* $< 0.05$).

**Table S10**: Linear $\beta$ and non-linear (quadratic and correlational, $\gamma$) selection gradients in **females** in **complex arenas**

| **Trait** | $\beta$ | $\gamma$ | | |
|---|---|---|---|---|
| | | **Beak length** | **Brain area** | **Gravid spot area** |
| $\boldsymbol{MS_M}$ | | | | |
| Beak length | -0.10 | -0.42 | 0.03 | 0.00 |
| Brain area | 0.34(.) | - | -0.01 | 0.00 |
| Gravid spot area | -0.04 | - | - | -0.21 |
| $\boldsymbol{MS_P}$ | | | | |
| Beak length | -0.10 | -0.33 | -0.14 | -0.06 |
| Brain area | 0.26(.) | - | 0.19 | 0.14 |
| Gravid spot area | 0.05 | - | - | -0.19 |

$\beta$ and $\gamma$ values are shown for both mating success characterization ($MS_M$ : number of matings; $MS_P$ : number of mates). Standard length was excluded from the model as multicollinearity was detected among explanatory traits. Significance effects are shown in boldface. Significance effects are shown in boldface (.: *p-value* $< 0.1$).

**Table S11**: The matrix of eigenvectors ($m$) and estimates of non-linear selection on the axes (eigenvalues, $\lambda$) described by the eigenvectors from the canonical analysis of gamma matrix $\gamma$ in **females** in **complex arenas**

| *m* | **Beak length** | **Brain area** | **Gravid spot area** | $\lambda$ | ***P-values*** |
|---|---|---|---|---|---|
| $\boldsymbol{MS_M}$ | | | | | |
| *m1* | -0.09 | -1.00 | 0.05 | 0.01 | 0.99 |
| *m2* | 0.04 | 0.04 | 1.00 | -0.20 | 0.34 |
| *m3* | 1.00 | -0.09 | -0.04 | -0.46 | 0.62 |
| $\boldsymbol{MS_P}$ | | | | | |
| *m1* | 0.21 | -0.94 | -0.28 | 0.27 | 0.72 |
| *m2* | -0.06 | -0.29 | 0.95 | -0.22 | 0.19 |
| *m3* | 0.98 | 0.18 | 0.11 | -0.40 | 0.14 |

$m$ values are shown for both mating success characterization ($MS_M$ : number of matings; $MS_P$ : number of mates). Standard length was excluded from the model as multicollinearity was detected among explanatory traits. Significance effects are shown in boldface. Significance effects are shown in boldface.

**Table S12**: The matrix of eigenvectors ($m$) and estimates of non-linear selection on the axes (eigenvalues, $\lambda$) described by the eigenvectors from the canonical analysis of gamma matrix $\gamma$ in **females** in **open arenas**

| $m$ | Beak length | Brain area | Gravid spot area | $\lambda$ | *P-values* |
|---|---|---|---|---|---|
| $MS_M$ | | | | | |
| *m1* | 0.72 | -0.08 | -0.69 | 0.62 | 0.47 |
| *m2* | 0.49 | 0.76 | 0.43 | -0.07 | 0.34 |
| *m3* | 0.49 | -0.65 | 0.58 | -0.79 | 0.15 |
| $MS_P$ | | | | | |
| *m1* | -0.57 | 0.02 | 0.82 | 0.50 | 0.21 |
| *m2* | 0.43 | 0.86 | 0.28 | -0.07 | 0.32 |
| *m3* | 0.70 | -0.51 | 0.50 | -0.49 | 0.10 |

$m$ values are shown for both mating success characterization ($MS_M$ : number of matings; $MS_P$ : number of mates). Standard length was excluded from the model as multicollinearity was detected among explanatory traits. Significance effects are shown in boldface. Significance effects are shown in boldface.

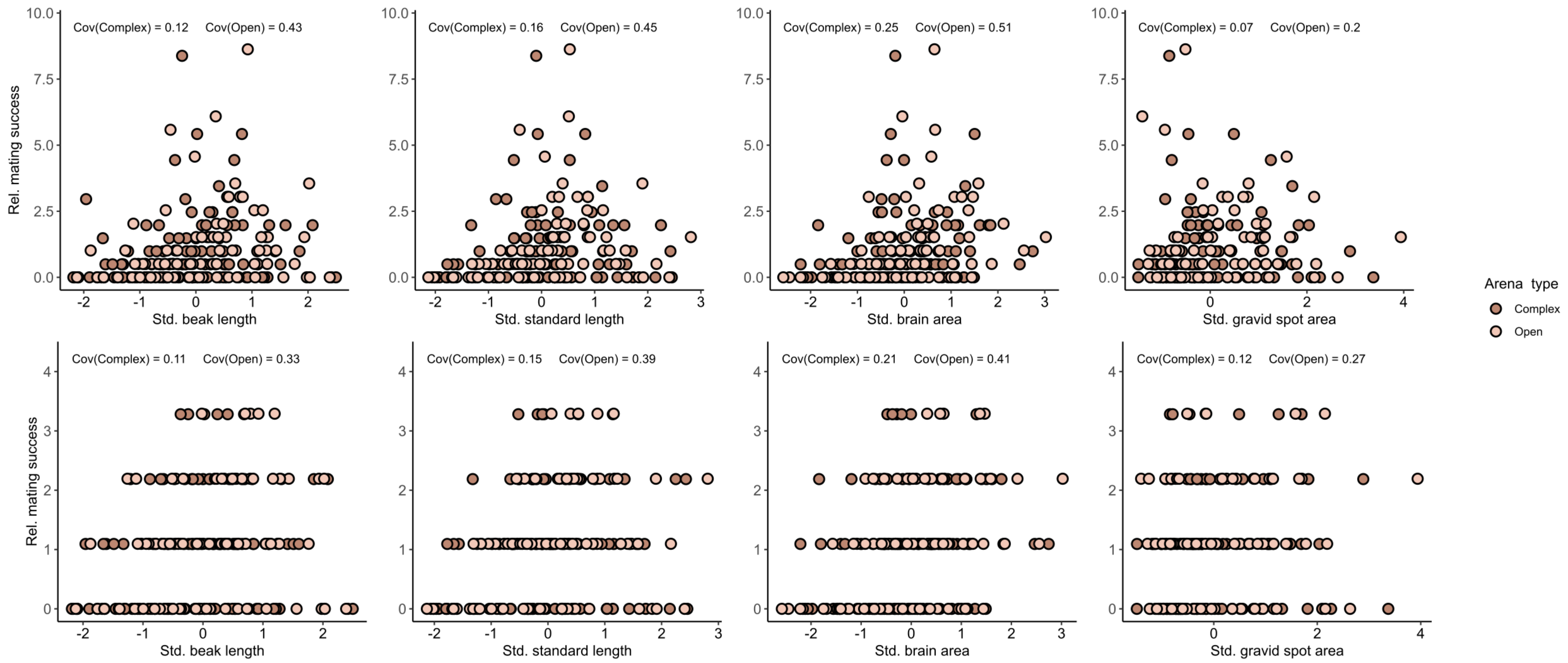


**Fig. S1**: Relationship between standardized traits and relative mating success success for females in open (light pink) and (dark pink) arenas. Covariance values obtained in each arena type are indicated above each plot.

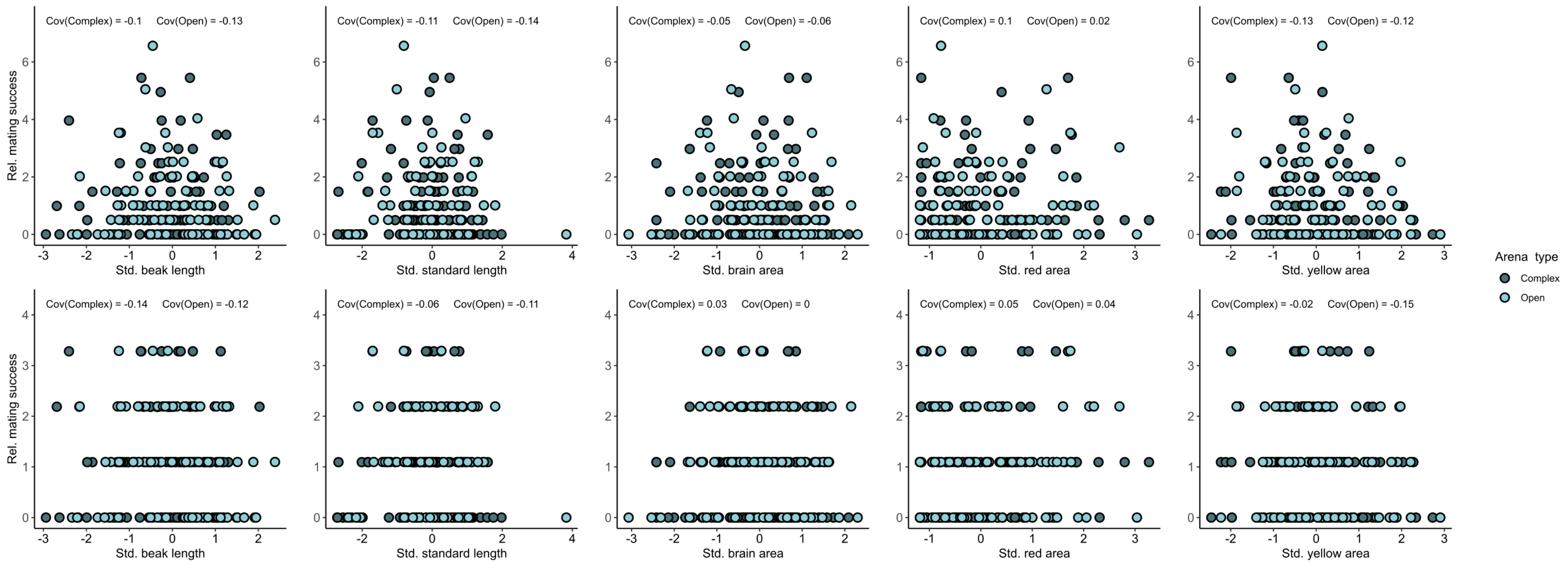


**Fig. S2**: Relationship between standardized traits and relative mating success success for males in open (light blue) and (dark blue) arenas. Covariance values obtained in each arena type are indicated above each plot.

**Table S13**: Influence of arena type on mating differentials (*m'*) on relative morphological traits

| Parameter | Mean complex (95% CI) | Mean open (95% CI) | Effect size (95% CI) | *P-values* |
|---|---|---|---|---|
| ***$MS_M$*** | | | | |
| **Females** | | | | |
| Beak length | -0.06 (-0.28,0.14) | 0.05 (-0.18,0.31) | -0.12 (-0.45,0.2) | 0.54 |
| Brain area | **0.21 (0.03,0.44)** | **0.24 (0.02,0.50)** | -0.03 (-0.35,0.30) | 0.90 |
| Gravid spot area | -0.03 (-0.29,0.21) | -0.10 (-0.46,0.23) | 0.07 (-0.34,0.50) | 0.70 |
| **Males** | | | | |
| Beak length | -0.03 (-0.25,0.18) | -0.03 (-0.21,0.16) | 0.00 (-0.29,0.28) | 1.00 |
| Brain area | 0.03 (-0.19,0.26) | 0.08 (-0.16,0.31) | -0.06 (-0.38, 0.28) | 0.75 |
| Total red areas | 0.11 (-0.14,0.37) | -0.01 (-0.25,0.24) | 0.12 (-0,24,0.47) | 0.51 |
| Total yellow areas | -0.09 (-0.38,0.14) | -0.06 (-0.26,0.14) | -0.03 (-0.38,0.27) | 0.85 |
| **ΔSex** | | | | |
| Beak length | 0.04 (-0.26,0.33) *p-value* = 0.85 | -0.08 (-0.39,-0.23) *p-value* = 0.66 | - | - |
| Brain area | -0.19 (-0.50,0.12) *p-value* = 0.30 | -0.16 (-0.51,0.18) *p-value* = 0.38 | - | - |
| ***$MS_P$*** | | | | |
| **Females** | | | | |
| Beak length | -0.05 (-0.23,0.13) | -0.04 (0.25,0.14) | 0.00 (-0.26,0.26) | 0.97 |
| Brain area | **0.17 (0.01,0.32)** | **0.16 (0.00,0.34)** | 0.01 (-0.23,0.26) | 0.95 |
| Gravid spot area | 0.04 (-0.14,0.21) | 0.04 (-0.16,0.26) | -0.01 (-0.28,0.27) | 0.96 |
| **Males** | | | | |
| Beak length | -0.14 (-0.34,0.07) | -0.05 (-0.23,0.14) | -0.08 (-0.36,0.19) | 0.53 |
| Brain area | 0.10 (-0.09,0.28) | 0.13 (-0.05,0.29) | -0.03 (-0.28, 0.22) | 0.82 |
| Total red areas | 0.06 (-0.12,0.25) | 0.02 (-0.17,0.23) | 0.03 (-0,24,0.30) | 0.81 |
| Total yellow areas | 0.02 (-0.18,0.19) | -0.12 (-0.30,0.06) | 0.13 (-0.13,0.38) | 0.33 |
| **ΔSex** | | | | |
| Beak length | -0.08 (-0.36,0.19) *p-value* = 0.54 | -0.004 (-0.26,0.26) *p-value* = 0.98 | - | - |
| Brain area | -0.07 (-0.31,0.18) *p-value* = 0.61 | -0.03 (-0.28,0.22) *p-value* = 0.81 | - | - |

To calculate relative morphological traits (e.g. brain size), we selected on the residuals from the regression of each trait (e.g. brain size) on body size. Confidence intervals in each arena type are obtained via bootstrap sampling and are shown for both sexes, for all traits and for both mating success characterisation ($MS_M$: number of matings; $MS_P$: number of mates). Sex difference observed within each arena type (ΔSex) are also represented. Significance of effects sizes is assessed via permutations tests (*P-values*). Significance effects are shown in boldface.
